# Spatial and developmental reprogramming enables root growth under high salinity

**DOI:** 10.64898/2026.09.15.751606

**Authors:** Thu T. Nguyen, Jason R. Garcia, Samadhi Wimalagunasekara, Richard S. Garcia, Hannah Holliday, Maria Iftesum, Elif Gediz Kocaoglan, James V. Moroney, Manas R. Gartia, José R. Dinneny, Maheshi Dassanayake

## Abstract

How do plant roots sustain growth under salinity levels that would normally arrest development? The extremophyte *Schrenkiella parvula* maintains root growth under high salinity through a previously unrecognized spatial reorganization of primary root development. The root elongation zone transitions into two distinct developmental states, the bulged and gap zones, during a transient growth pause that precedes renewed tip growth and localized lateral root emergence. In Arabidopsis, a related salt-sensitive species, the cells in the elongation zone simply burst. Jasmonic acid was found to be necessary to initiate this transition in *S. parvula*, while spatially coordinated jasmonic acid and auxin accumulation accompanies zone establishment and maintenance. We constructed zone-resolved transcriptomic networks to distinguish developmental from salt-responsive processes and identified their interactions under salinity stress. Salt-induced zone-specific networks revealed distinct regulatory programs coordinating hormone signaling, cell wall remodeling, and water, ion, and reactive oxygen species homeostasis. Cell layer-specific Raman spectroscopy, UHPLC-MS/MS metabolite profiling, and cellular imaging provide independent support for the transcriptomic networks, linking spatially organized cellular responses with salt stress mitigation, suppression of cell death, and continued growth. We also identified orthologous genes potentially recruited into novel regulatory roles that support growth under stress. Together, these findings establish spatial developmental reprogramming as a strategy for sustaining root growth under salinity and identify candidate mechanisms for improving crop salt tolerance.

## INTRODUCTION

High salinity presents one of the most formidable challenges to plant cell function, simultaneously disrupting osmotic, ionic, oxidative, and mechanical balance. Because the root is the first organ to encounter salt in the soil, root cells experience the earliest and most severe effects of salt stress across structural, physiological, and molecular levels. Most land plants have structural barriers in the root that prevent water loss and damage from excess salts. However, at the root apex (the meristematic and elongation zones), cells are actively dividing, expanding and differentiating, before protective barriers have fully matured, making this region particularly vulnerable to salt-induced damage. As Na⁺ enters the root, it weakens the pectin matrix of the cell wall, disrupts ion homeostasis by promoting K⁺ loss, and alters membrane permeability, thereby compromising both extracellular and intracellular structural stability. At moderate salinity (140 mM NaCl in *Arabidopsis thaliana*), the receptor-like kinase FERONIA senses salt-induced mechanical stress of the cell wall and activates localized Ca²⁺ signaling to preserve cell wall integrity. When this protective pathway is overwhelmed or disrupted under higher salinity, cortical and epidermal cells in the elongation zone undergo abnormal swelling and ultimately rupture under turgor pressure, arresting primary root growth. Concurrently, intracellular Na⁺ accumulation induces oxidative stress through excessive production of reactive oxygen species (ROS), causing damage to DNA, proteins, and membranes, disrupting cellular metabolism, and ultimately activating programmed cell death ^1^. During prolonged salt stress, these interconnected stresses initiate a cascade of cellular and physiological dysfunction that irreversibly impairs root function, triggers systemic stress responses, and ultimately leads to whole-plant death under high salinity ^2,3^.

Unlike the vast majority of land plants, which are highly susceptible to salinity, many halophytic extremophytes have evolved the ability to sustain root growth in saline soils. This exceptional resilience arises from coordinated physiological and molecular adaptations that preserve growth despite prolonged exposure to high salt ^4,5^. Although numerous components of salt tolerance have been identified, how protective mechanisms are coordinated in the most vulnerable root tissues to stabilize cellular structure, minimize salt-induced damage, and sustain growth over time, remain largely unresolved in model extremophytes. Deciphering how salt-adapted plants prevent the progression from early stress response to irreversible cellular damage is therefore a major knowledge gap in plant stress biology and is essential for understanding the mechanisms that facilitate salt tolerance.

In this study we used the salt-adapted extremophyte model, *Schrenkiella parvula (S. parvula)* to examine how vulnerable root tissues in the root tip are stabilized and recover growth under high salinity. Previous studies have shown that *S. parvula* can sustain growth and cellular function under salt concentrations that severely inhibit *A. thaliana.* Salinity levels of 200 mM NaCl are generally considered severely inhibitory to crop growth, yet the capacity of some plants to sustain or recover root growth at this salinity remains poorly understood ^6–9^. How spatiotemporal reprogramming across primary root tip developmental zones enable extremophytes to establish and maintain a new developmental equilibrium under salt stress remains unknown. Here we combined spatially resolved physiological, transcriptomic, metabolite, and developmental analyses in *S. parvula* to uncover how vulnerable root tissues are protected under high salinity. We identify an adaptive program that enables *S. parvula* to spatiotemporally reorganize root developmental zones, restore structural stability, suppress oxidative stress, and prevent irreversible tissue failure.

## RESULTS

### Dose-dependent salinity thresholds define developmental tipping points in primary root growth of *S. parvula*

The impacts of salt stress are most severe during the seedling stage ^8,10,11^. To determine the salinity threshold at which prolonged salt exposure begins to limit primary root growth, we grew *S. parvula* seedlings for one week on standard growth medium supplemented with 0 to 400 mM NaCl (Figures 1A, B). Primary root growth enhanced at moderate salinities but declined progressively as salinity increased, with a clear growth penalty emerging at ≥200 mM NaCl and complete inhibition of primary root tip growth at ≥250 mM NaCl. Based on these responses, we define high salt stress as 200 mM NaCl and extreme salt stress as ≥250 mM NaCl throughout this study for *S. parvula*. Notably, at 200 mM NaCl, the primary root growth was transiently paused during the first day of exposure before resuming continuous growth (Fig. 1B). These results reveal a dose-dependent developmental transition in *S. parvula* primary roots, from growth enhancement under moderate salinity to growth inhibition and eventual root tip arrest under extreme salinity.

**Figure 1.**
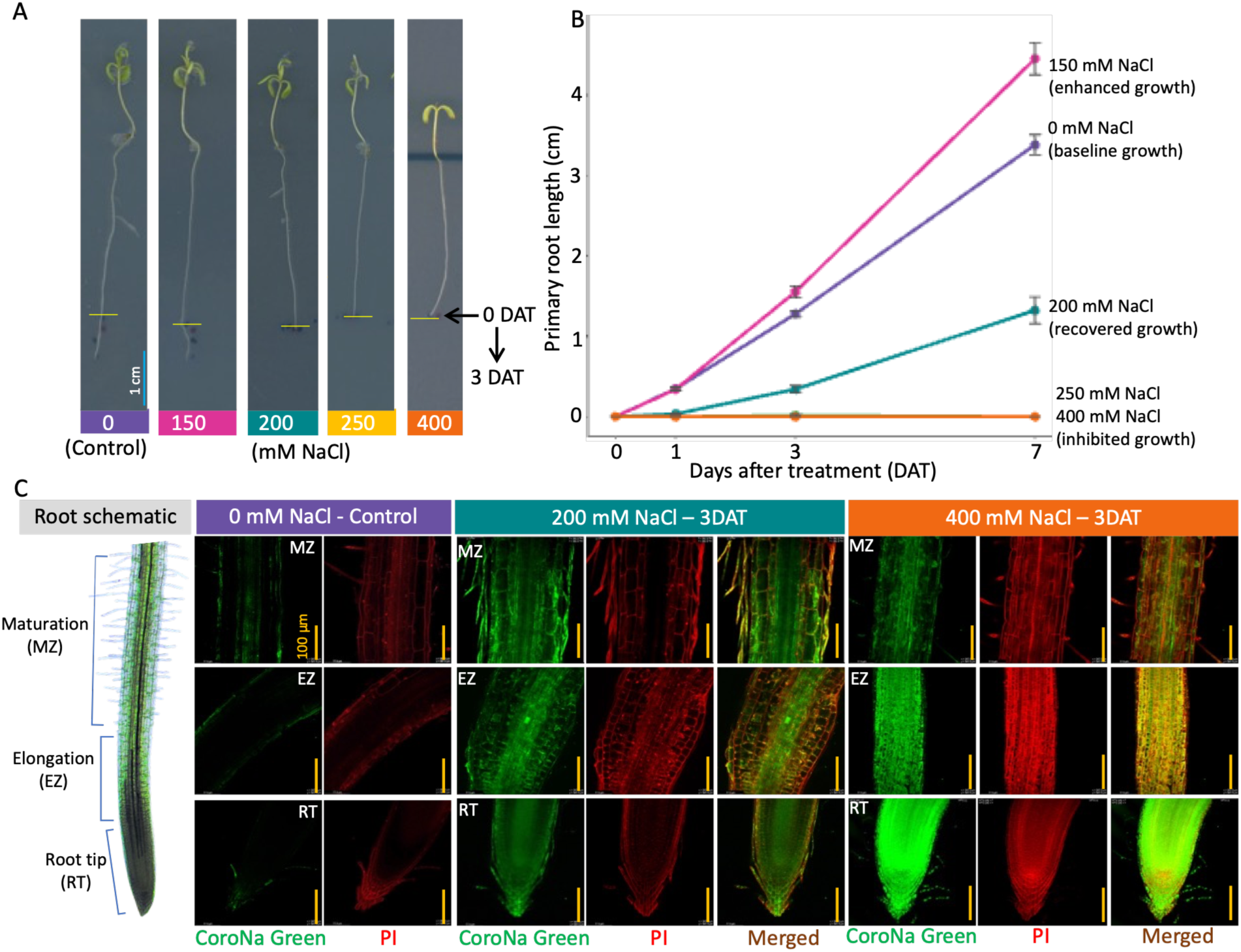
*Schrenkiella parvula* adjusts primary root growth and spatially restricts cell death under increasing salinity. **(A)** Ten-day-old seedlings transferred to 1/4^th^ MS medium supplemented with increasing salt concentrations and imaged at three days after treatment (DAT). The yellow lines indicate the tip of the root on Day 0 of transfer. **(B)** Primary root growth under the conditions examined in (A) transitioned from enhanced growth at 150 mM NaCl, to transient growth arrest followed by recovery at 200 mM NaCl, and to complete growth arrest at ≥250 mM NaCl, relative to the control over the seven-day observation period. Error bars indicate standard error. Non-overlapping data points at 3 and 7 DAT differ significantly from the control (*p < 0.05*, Student’s t-test). **(C)** Sodium accumulation in the root tip (RT), elongation zone (EZ), and the maturation zone (MZ), increases with external salinity as indicated by CoroNa Green bound to Na⁺. Propidium iodide (PI) stains cell walls but exhibits stronger fluorescence in membrane compromised cells after additional binding to nucleic acids. As salinity increases, PI staining progressively expands from the root tip into the elongation zone, indicating widespread loss of membrane integrity.

We next examined how sodium accumulation and cell viability varied across root developmental zones with increasing salinity. Under high salt stress, CoroNa Green staining revealed elevated sodium accumulation throughout the root compared to control seedlings grown without added NaCl (Fig. 1C). Sodium was preferentially localized to the stele from the root tip through the maturation zone, whereas lower levels accumulated in the surrounding cell layers between the epidermis and pericycle. Despite this increase in sodium, cells in the meristematic and elongation zones remained viable, as indicated by propidium iodide staining restricted to the cell walls (Fig. 1C, Figure S1). In contrast, under extreme salt stress, sodium accumulated uniformly across all cell layers from the epidermis to the stele within the root tip and elongation zone, coinciding with widespread intracellular propidium iodide staining indicative of loss of cell viability. By comparison, cells in the maturation zone remained largely viable, with sodium continuing to be preferentially confined to the stele (Fig. 1C).

Together, these results indicate two developmental tipping points in *S. parvula* primary roots. The first tipping point is the transition from growth enhancement at moderate salinity to a reduced but recoverable growth state at high salinity. The second tipping point is the transition from high salt to extreme salinity, when root tip death inhibits primary root growth, which is subsequently restored through lateral root development in the maturation zone. However, at high salinity (the transition between the two tipping points), the plant restores primary root growth by maintaining root tip integrity, whereas at extreme salinity, irreversible damage to the root tip terminates apical growth. In this study, we focused on the adaptive mechanisms underlying growth re-establishment through a new developmental equilibrium at high salinity, corresponding to the transition point.

### Ionic stress induces two new developmental zones replacing the elongation zone in *S. parvula*

Epidermal and cortical cells in the early elongation zone underwent pronounced radial expansion under moderate and high salinity, forming a characteristic bulged zone (Figure 2A, Supplementary Video S2). Although the bulged zone formed consistently, its morphology varied among seedlings (Figure S2). In contrast to the bulged zone, cells proximal to the maturation zone remained unexpanded, forming a distinct root hair-free region that we refer to as the gap zone (Fig. 2A, Supplementary Video S2). These zones developed in more than 75% of seedlings exposed to 150–200 mM NaCl but were absent at lower salinities (Fig. 2A, B, Supplementary Video S1). Under extreme salinity, however, cells in the early elongation zone burst, resulting in the loss of root tip structural integrity (Fig. 2A, B, Supplementary Video S3). Once initiated, these developmental changes were fully established within 24 h of salt exposure, coinciding with the transient pause in primary root growth before growth resumed under high salinity (Fig. 1B, Supplementary Video S2). Cells within the bulged zone were both significantly shorter and wider than elongation zone cells under control conditions, although cell number remained unchanged (Fig. 2C). Therefore, the bulged zone is formed by radial cell expansion, not by increasing the number of cell or cell layers.

**Figure 2.**
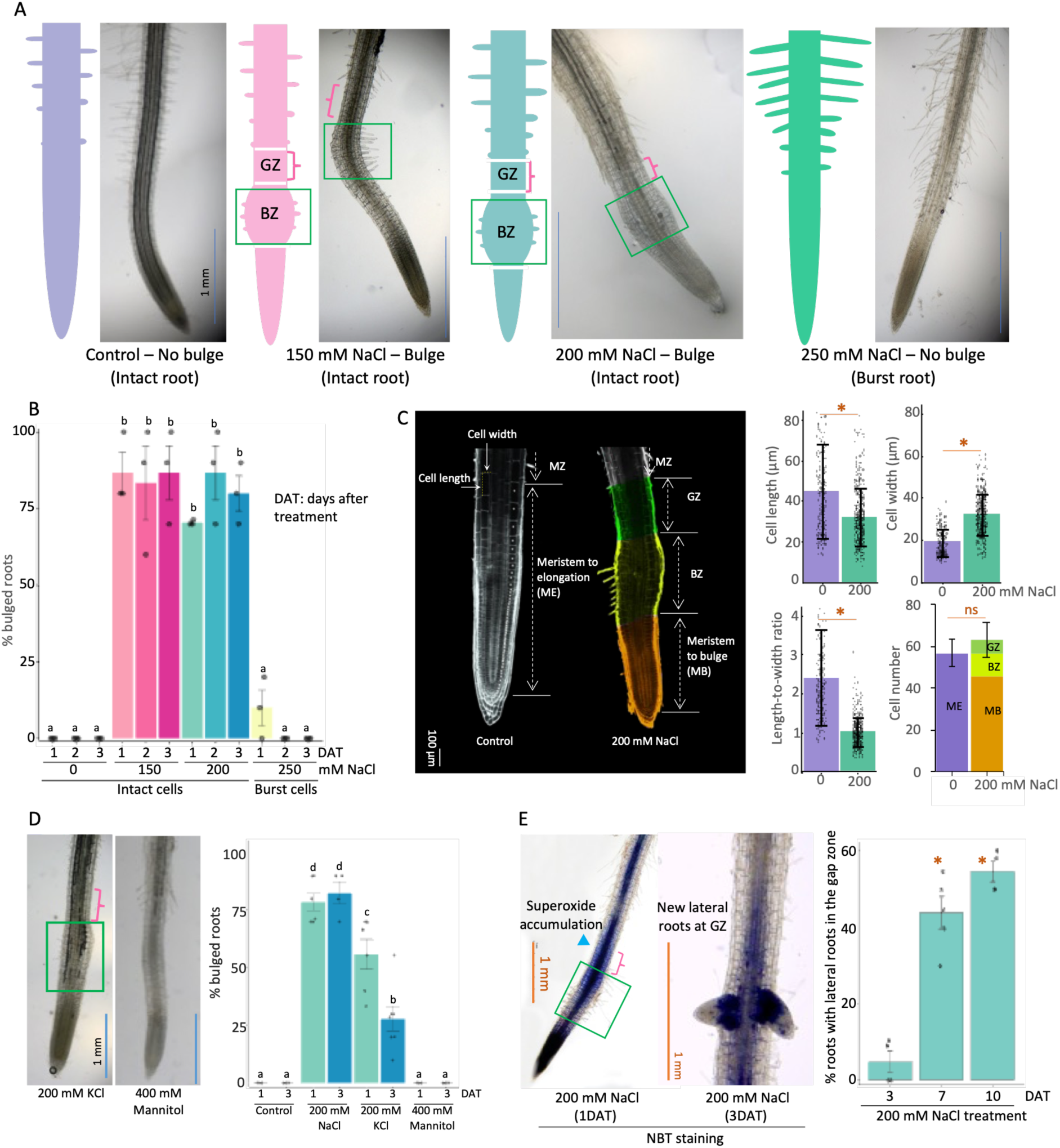
Ionic stress induces two new developmental zones - the bulged and gap zones, in *S. parvula* primary roots under increasing salinity. **(A)** Structural changes of primary roots in response to increasing salt. **(B)** Percentage of roots exhibiting bulge formation under salt treatment. **(C)** Grayscale confocal micrographs of the root apex were post-processed with digital coloring to highlight developmental zones under 200 mM NaCl. Cell length and width measurements under control conditions were obtained from cells spanning the meristem to the elongation zone (ME), whereas measurements under salt treatment included cells from the meristem through the bulged and gap zones (MB+BZ+GZ). The maturation zone (MZ) was marked by the appearance of the first root hair at its distal boundary. Representative cortical cells counted in the analysis are indicated by a white asterisk in the control root. **(D)** Bulge formation was induced by 200 mM KCl but not by 400 mM mannitol, despite equivalent osmotic stress. **(E)** NBT staining reveals that superoxide accumulates within 1 day of salt treatment, preceding lateral root initiation in the gap zone by day 3. The squares indicate the bulged zone, and the brackets indicate the gap zone in panels A, D, and E. Treatments were repeated at least three times, with each biological replicate including at least 10 seedlings for panels B-E. Error bars indicate standard error. Statistical analyses for these were performed using one-way ANOVA followed by Tukey’s post hoc test (B, D) and Student’s t-test (C, E); (*p_adj_ < 0.05*); significant differences among groups are shown by different lowercase letters or an asterisk. DAT, days after treatment.

The developmental transition into the bulged and gap zones under moderate to high salt was induced by ionic stress. Mannitol was used to distinguish osmotic from ionic effects and did not induce this response, whereas both Na⁺ and K⁺ treatments did (Fig. 2D). Concurrently, the gap zone accumulated superoxide within one day of salt treatment, preceding lateral root initiation from this region over time (Fig. 2E). These findings suggest that elongation-zone remodeling under high salinity may represent either an adaptive process that facilitates root growth recovery or an early stage of developmental failure preceding recovery, thereby providing a developmental framework for investigating the mechanisms that restore root growth.

### Spatially and temporally coordinated jasmonic acid-auxin synergy mediates salt-induced developmental transition

Elevated levels of ABA, JA, and auxin are well documented to suppress primary root growth in Arabidopsis ^12,13^. Because we observed the opposite growth response in *S. parvula*, we first sought to determine how these hormones may be differently organized in this extremophyte. We found that exogenous application of jasmonic acid (JA; applied as methyl jasmonate) induced the developmental transition of the elongation zone into the bulged and gap zones in the absence of salt, and this response was not further enhanced by salt treatment (Figures 3A, B). The JA biosynthesis inhibitor, sodium diethyldithiocarbamate trihydrate (DIECA) prevented this developmental transition under salt stress (Fig. 3A, B). In contrast, exogenous auxin (applied as IAA), promoted root hair development within the bulged zone but did not induce the bulge formation without salt (Fig. 3C, S3). Consistent with this, the auxin signaling inhibitor Auxinole did not prevent the salt-induced bulge formation. Additionally, exogenous ABA application did not cause root bulging in the absence of salt (Supplementary Video S4). Together, these results indicate that JA mediates the salt-induced developmental remodeling of the elongation zone into the bulged and gap zones and that the endogenous JA accumulation under high salinity is sufficient to initiate this process.

**Figure 3.**
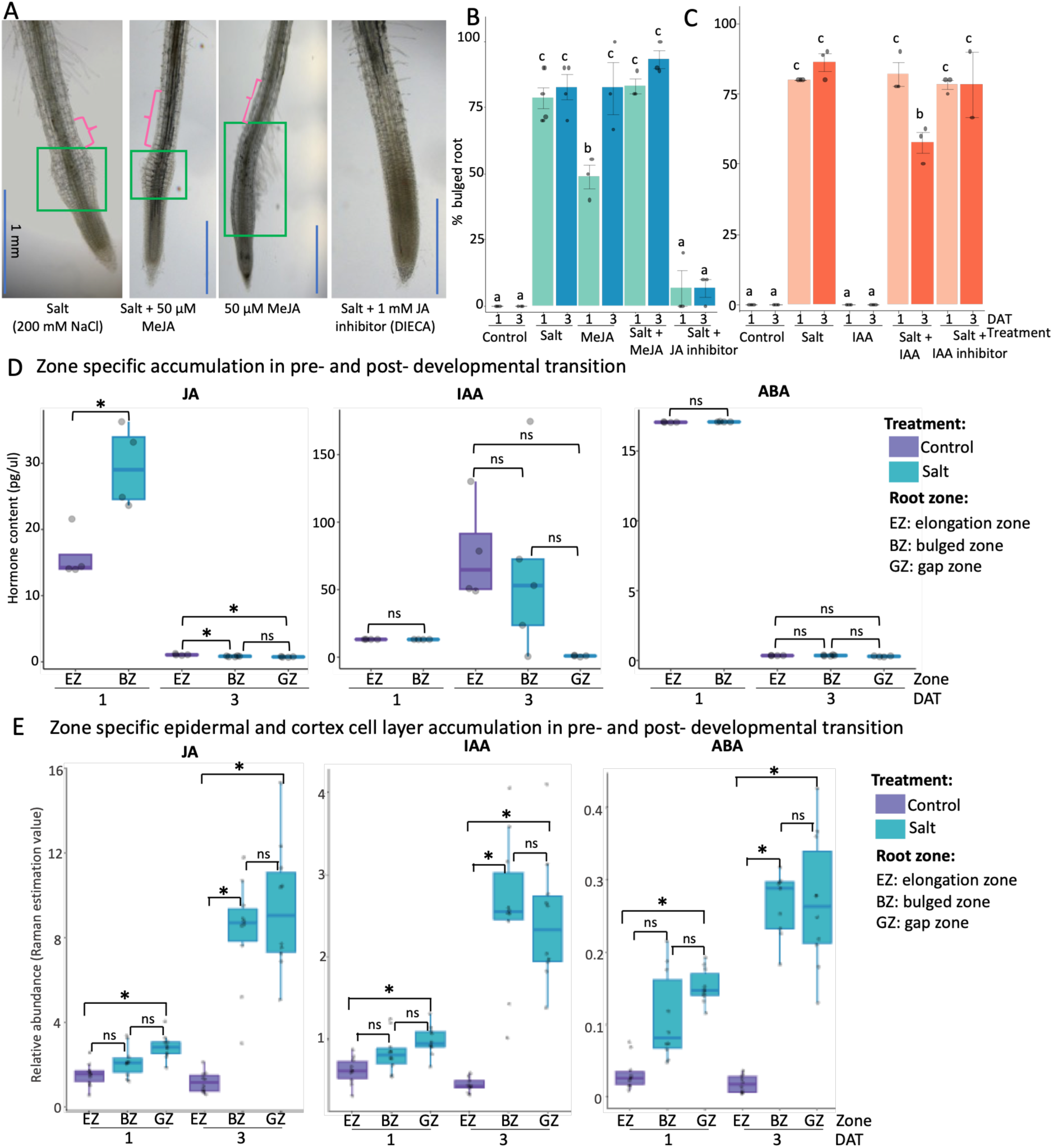
Spatiotemporal hormone accumulation under salt stress is associated with gap and bulged zone formation in *S. parvula* roots. **(A)** Jasmonic acid (JA) induces bulge formation with or without salt, whereas its biosynthesis inhibitor, DIECA suppresses bulge formation under salt. The squares indicate the bulged zone, and the brackets indicate the gap zone. **(B)** The proportion of roots exhibiting bulge formation under salt treatment was comparable to that induced by exogenous methyl jasmonate (MeJA) in the absence or presence of salt. Inhibition of endogenous jasmonic acid biosynthesis by DIECA completely suppressed bulge formation. **(C)** Exogenous IAA alone did not induce bulge formation compared to the control. IAA did not increase the proportion of bulged roots under salt, and inhibition of endogenous auxin signaling with auxinole had no effect on salt-induced bulge formation. **(D)** Whole-zone tissue sections (containing multiple cell layers) and **(E)** *in situ* epidermal and cortical cell layers were used from the elongation zone (EZ) under control conditions and the bulged (BZ) and gap (GZ) zones following 200 mM NaCl treatment on Day 1 (growth paused under salt) and Day 3 (growth resumed under salt) for quantification of endogenous JA, IAA, and ABA. Statistical analyses for these were performed using one-way ANOVA followed by Tukey’s post hoc test (B, C) and using Welch’s two-sample *t*-test followed by Holm correction to adjust for multiple testing (D, E); (*p_adj_ < 0.05*). Outliers were removed using Grubbs’ test (*p* < 0.05) (D, E). Error bars indicate standard error, and boxes indicate the interquartile range with median. Significant differences among groups are shown by different lowercase letters or an asterisk. Same letter or ns-not significant; DAT-days after treatment.

To determine whether JA differentially accumulates during this developmental transition, we quantified hormone levels in each developmental zone over a 3-day period. When compared between the control elongation and salt-treated bulged zones, of the three hormones examined, only JA showed a significant increase, whereas auxin (IAA) and abscisic acid (ABA) remained unchanged (Fig. 3D). JA accumulated significantly in the bulged zone within one day of salt treatment, coinciding with the time taken to complete the bulge formation (Fig. 3D, 2B). By day 3, JA levels had declined in the newly established developmental zones while the root had resumed growth (Fig. 3D, 1B), although they remained higher than in the elongation zone, indicating that JA accumulation is temporally aligned with the developmental transition (Fig. 3D).

Given that auxin promoted root hair development within the bulged zone (Fig. S3) and ABA responses are cell type-specific in *S. parvula* roots ^14^, the absence of changes in their overall accumulation (Fig. 3D) suggests that zone-level hormone measurements alone may be insufficient to explain the distinct developmental identities of the bulged and gap zones. Instead, these phenotypes may depend on temporal and cell type-specific distribution, signaling, or sensitivity. Therefore, we examined whether cell layer-specific hormone distributions could further explain the distinct cellular phenotypes of the bulged and gap zones. To achieve this spatial and temporal resolution, we used Raman spectroscopy ^15–17^ focusing the laser’s *in situ* probing depth to span the epidermis and cortex cell layers at each zone. We found that JA, IAA, and ABA accumulated to significantly higher levels in the outer cell layers of both the bulged and gap zones than in the elongation zone, showing distinct spatiotemporal patterns (Fig. 3E). On day 1, when the developmental transition is established, IAA accumulated significantly in the gap zone relative to the elongation zone, but not in the bulged zone (Fig. 3E). Lower auxin levels are known to permit cell expansion, whereas higher levels increase cell wall rigidity and inhibit cell expansion ^18^. This may explain why outer cells remain unexpanded in the gap zone but expand in the bulged zone. By day 3, a subsequent auxin increases together with higher JA and ABA in both the bulged and gap zones. This may reflect an early stage of developmental failure. Alternatively, these changes likely stabilize these developmental regions, preventing excessive, destructive expansion under high salinity. Collectively, our findings suggest that localized JA accumulation mediates the initial developmental transition of the elongation zone under salt stress, whereas differential auxin accumulation drives the distinct cell expansion patterns separating the bulged and gap zones.

### Spatial organization of transcriptional networks shapes distinct developmental states during salt stress

We posited that salt-induced structural changes in the root are driven by underlying transcriptional reprogramming. To capture these transcriptional dynamics, we examined *S. parvula* seedlings exposed to 200 mM NaCl for three days; at this stage, the bulged and gap zones were fully established, and primary root growth had resumed, reflecting a new growth– stress equilibrium (Fig. 1B, 2B). We sectioned the primary roots into distinct regions along the root axis. Under control conditions, roots were divided into three zones: the root tip/meristematic zone (RT), elongation zone (EZ), and maturation zone (MZ). Under salt, roots were divided into four zones: the root tip (RT), bulged zone (BZ), gap zone (GZ), and maturation zone (MZ). Notably, for both control and salt-treated MZ samples, we collected only ∼1 cm above the EZ (control) or GZ (salt). This sampling design excluded older tissue that had initiated secondary growth or lateral root development. By avoiding these additional structural layers in the mature roots, we eliminated confounding transcriptional signatures to isolate the developmental reprogramming associated with bulge formation under salt stress.

Transcriptomic profiling revealed that control and salt-treated primary roots formed distinct clusters, reflecting extensive transcriptional reprogramming under salt stress (Figure 4A, Table S1). The bulged and gap zones each exhibited transcriptional states distinct from one another and from the elongation zone. A comparison of cumulative gene expression distributions between control and salt-treated conditions revealed that transcript abundance was predominantly reduced under salt stress (Fig. 4B, Table S1).

**Figure 4.**
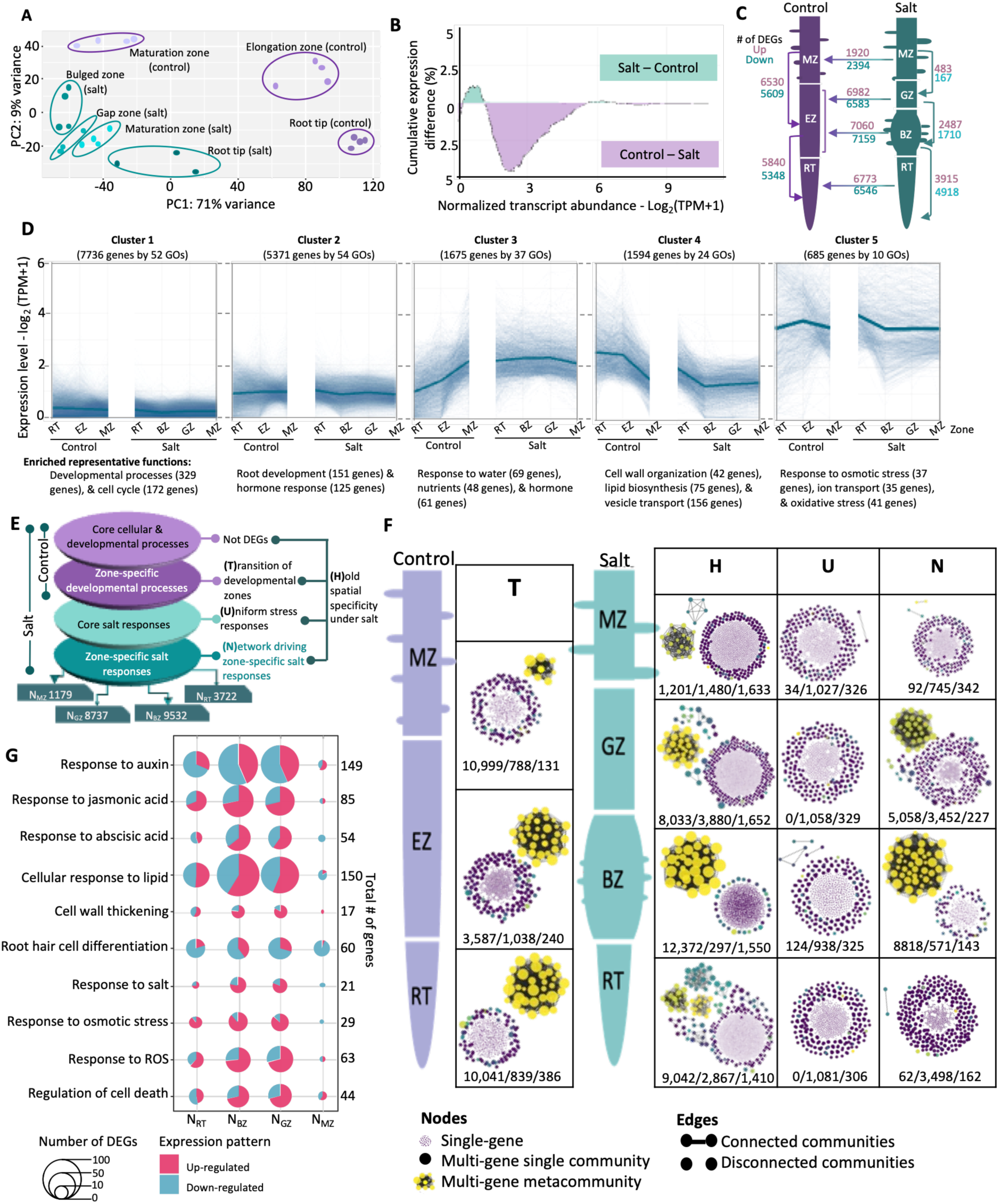
Layered and connected transcriptional networks coordinate spatial and developmental adaptation to salt stress. **(A)** Principal component analysis (PCA) of the 2,000 most variable genes. Transcriptome profiles from 3 - 4 biological replicates for each developmental zone formed distinct clusters. **(B)** Cumulative gene expression distribution analysis comparing control and 200 mM NaCl treatments. The distribution reveals a global reduction in transcript abundance under salt stress, while a distinct subset of low-expressed genes (<2 TPM) was induced. **(C)** Number of DEGs during developmental transitions under control conditions (left) and developmental transitions under salt stress (right). Differentially expressed genes (DEGs) were identified for each pairwise comparison indicated by unidirectional arrow between developmental zones and treatments (*p_adj_* ≤ 0.05). 3 - 4 biological replicates (≥100 plants per replicate). The numbers of upregulated (lilac) and downregulated (teal) DEGs are indicated adjacent to the arrows, which denote salt/control or older/younger tissue comparisons. MZ, maturation zone; EZ, elongation zone; BZ, bulged zone; GZ, gap zone; RT, root tip. **(D)** Co-expression gene clusters identified by k-means clustering for all DEGs among comparisons across developmental zones under control and salt treatments. Each cluster was analyzed for functional enrichment, with representative enriched biological processes shown below each cluster. Functional enrichment was based on *S. parvula* gene annotations, *p_adj_* ≤ 0.05. **(E)** Analysis framework illustrating the layered transcriptional networks that shape root development and salt responses. Baseline root development is supported by core cellular and developmental processes in addition to zone specific zone-specific programs associated with developmental transitions (T). Salt stress adds a uniform response shared across root zones (U) and distinct transcriptional networks that drive zone-specific salt responses (N), while H represents transcriptional programs that retain spatial specificity under salt stress. The number of DEGs for the N networks are listed. **(F)** Networks representing the complexity of transcriptional responses from regular development to salt-induced developmental changes. Genes in T, H, U, and N networks were clustered using Louvain clustering based on pair-wise Pearson correlation between genes within each set. Gene pairs with |r| ≥ 0.90 and *p ≤* 0.001 were selected to represent communities with edges between nodes. The numbers under each network show the number of DEGs in the order for multi-gene metacommunity, multi-gene single community, and single gene clusters respectively. The color gradient of each circle represents each community. **(G)** Enriched functional processes (*p*-values ≤ 0.05) represented by N networks.

To identify the transcriptomic shifts driving root architectural remodeling, we mapped differentially expressed gene (DEG) patterns across developmental zones under control and salt stress conditions to trace the transitions between distinct transcriptomic states (Fig. 4C, Table S3). The transition from the EZ to the salt-induced BZ and GZ represented the greatest divergence (based on the number of DEGs) in the spatial transcriptomic landscape. In contrast, the MZ undergoes a minimal transcriptomic readjustment by producing the fewest DEGs during the control-to-salt transition (Fig. 4C, Table S3).

To determine if functionally related genes define spatially organized transcriptomes, we examined co-expression profiles across the root developmental axis and tested which functional processes are enriched in each zone (Fig. 4D, Table S4). To manage developmentally coordinated zones formed under salt stress, *S. parvula* activates transcriptional responses by constitutively elevating water, ion transport, hormone-regulation, osmotic, and oxidative stress defenses (Cluster 3 and 5), while down-regulating cell wall and membrane organization genes (Cluster 4) to structurally accommodate morphological adjustments. Our results demonstrate that the anatomical emergence of the bulged and gap zones is accompanied by discrete, zone-specific transcriptional states represented by coordinated functional modules, shifting our understanding of root stress responses in *S. parvula* from a uniform tissue-wide reaction to a highly compartmentalized developmental adaptation.

We aimed to isolate the transcriptional networks that mediate salt tolerant responses to reach the new developmental equilibrium. We anticipated that these dynamics would be driven by DEGs; however, individual DEGs that serve as conditional marker genes or are unique to a single developmental zone would be insufficient to capture the salt-induced transcriptional network driving spatially restricted cellular processes. Instead, we reasoned that zone-specific outcomes depend on distinct transcriptional layers—coordinated networks of co-expressed modules that regulate collections of functional processes rather than single, isolated genes (Fig. 4E). Therefore, we evaluated transcriptomic profiles across developmental zones under both control and salt conditions to define four modules (Fig. 4E, S4A) (see Methods):

1. **T** (Transition of developmental zones): this set acts as a transcriptionally coordinated module to govern developmental shifts along the root axis in the control condition.
2. **H** (Hold spatial specificity while influenced by salt): this set is affected by salt and developmental processes that operate independently of or in addition to core salt responses.
3. **U** (Uniform stress responses): this set serves as a core, salt-responsive module acting across cells, tissues, and developmental zones.
4. **N** (Network driving spatially restricted stress responses): this set governs localized salt responses that are restricted to each root zone.

By analyzing the DEGs detected within each individual zone, we deduced the specific gene sets for each of the T, H, U, and N modules (Fig. S4B-F, Table S5) (see Methods). We expected the N modules to show the highest spatial specificity (Fig. 4E). The N_GZ_ (gap zone) and N_BZ_ (bulged zone) networks formed the largest modules, with the greatest number of differentially expressed genes (Fig. 4E, S4F, Table S5).

Under salt stress, the baseline control network (T) reorganizes into three layered network architectures (H, U, and N), characterized by different levels of connectivity and coordinated expression among subnetworks (represented as co-expressed gene communities in Fig. 4F, Table S6). Salt-induced genes in the H networks form larger, multi-gene metacommunities than those in the U networks, which represent broadly responsive salt-induced genes shared across all root zones and are organized into smaller, more independently regulated communities. In contrast, the zone-specific salt-induced genes comprising the N networks exhibit the highest degree of network organization, forming dense, highly connected metacommunities with the strongest coordination among gene networks in the bulged and gap zones. Thus, the N networks minimize isolated single genes while maximizing coordinated expression across the network (Fig. 4F, Table S6).

To elucidate the primary pathways behind this localized developmental reorganization, we investigated the N networks as the principal transcriptional framework governing these responses under salinity. While broad cellular processes were shared across all N networks, hormone signaling, osmotic and ROS responses, and cellular structural modification pathways comprised a larger proportion of the N_GZ_ and N_BZ_ networks (Fig. 4G). Contrastingly, the N_MZ_ module displayed minimal transcriptional responses across most categories. Shared functional pathways showed higher transcriptional similarity between N_GZ_ and N_BZ_ networks than with either of the remaining zones (Fig. 4G, Table S7). This similarity may reflect the common origin of the bulged and gap zones from the elongation zone before salt exposure. On one hand, these results suggest that N_GZ_ and N_BZ_ share a substantial transcriptional network underlying their similar functional composition. On the other hand, their morphological and molecular phenotypes are distinct (Fig. 2 and 4), suggesting that these shared networks are differentially deployed to establish zone-specific processes. We therefore aimed to identify the shared and unique genes within the N networks that collectively support cellular processes required for stress adaptation and resumed growth in both zones, while also determining how divergent regulation of shared genes contributes to the distinct functional programs of the bulged and gap zones.

### Differential deployment of shared networks underlies bulged and gap zone identities

To evaluate how gene expression is partitioned between the bulged and gap zones, we compared the N_BZ_ and N_GZ_ networks and examined the proportional contributions of zone-specific genes to shared functional processes (Fig. 5A, Table S8). Our goal was to identify processes that are prioritized within coordinated yet spatially restricted programs, which may be obscured when each zone is analyzed independently. Uniquely expressed genes in both zones contributed similarly to ROS scavenging and redox balance under high salinity. However, there was a greater contribution of genes associated with salt and water stress, cell growth, lipid metabolism, and transport from N_BZ_ suggesting a greater reliance on these processes in the bulged zone than in the gap zone. Conversely, cell wall organization was more strongly represented in N_GZ_ (Fig. 5A, Table S8). Additionally, the shared genes between the two networks were predominantly enriched in responses to water and salt, lipid metabolism, root morphogenesis, programmed cell death, and cell wall thickening (Fig. 5A). Because these processes correspond to physiological adaptations required to withstand osmotic and ionic stress, their enrichment suggests that they provide a common functional foundation for maintaining cellular integrity and enable growth recovery in both zones, upon which zone-specific transcriptional programs are layered (Fig. 5A, Table S8).

**Figure 5.**
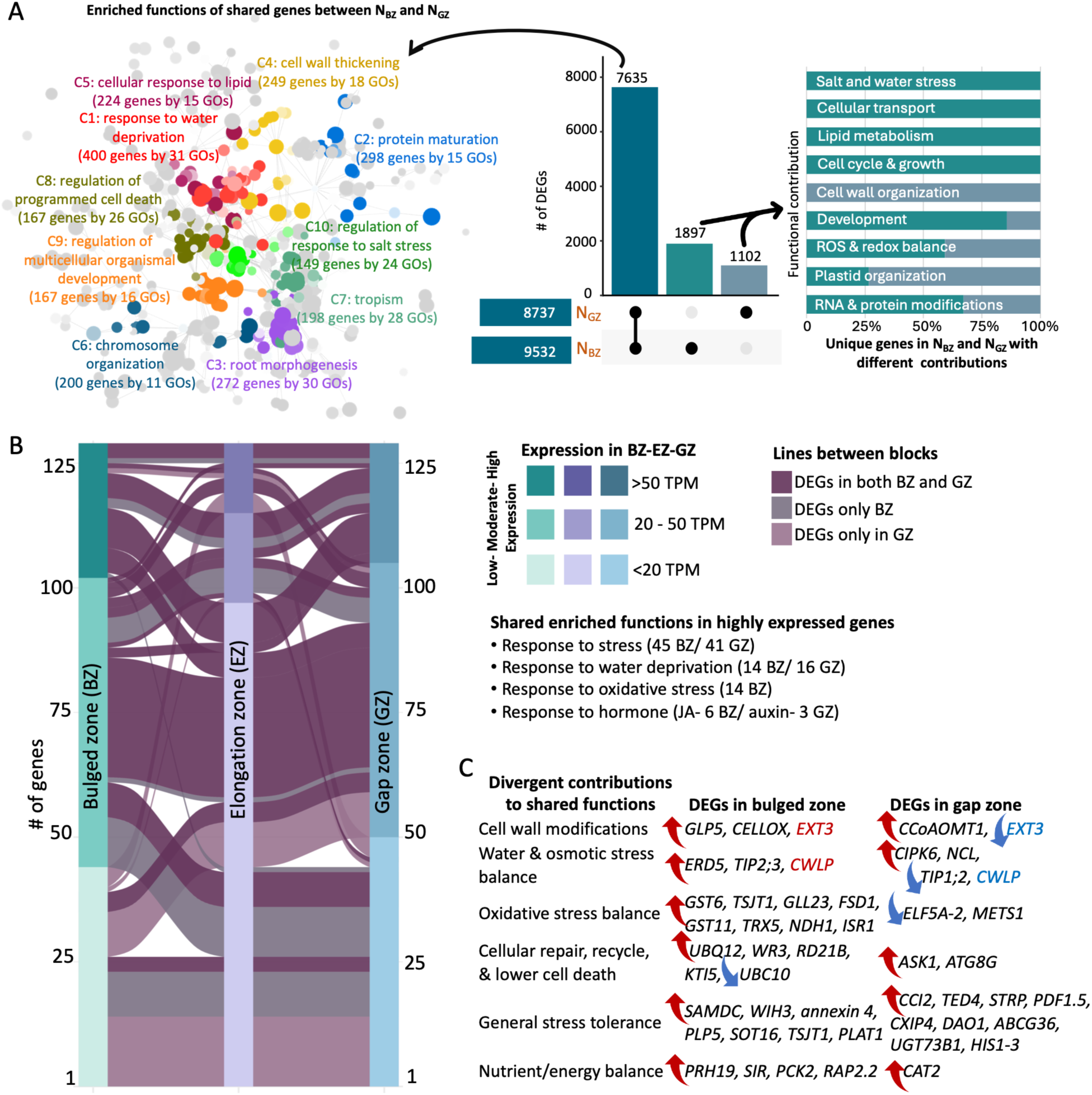
Bulged and gap zones share core salt-responsive programs but prioritize distinct functional processes in the salt-induced zone-specific transcriptional networks. **(A)** The shared and distinct DEGs in the N_BZ_ and N_GZ_ networks. Bars indicate the number of genes shared between bulged and gap zones (= connected dots) as well as those uniquely expressed in each zone (= single dots), illustrating both shared and unique transcriptional responses to salt stress. Shared processes contributed by uniquely expressed genes in the two zones are shown as a stacked horizontal bar chart on the right. Enriched functions among shared DEGs between the N_BZ_ and N_GZ_ networks on the left. Nodes represent GO terms, scaled by gene number. Colors group functionally related terms, with darker shades indicating stronger enrichment (lower adjusted *p*-values). Edges represent gene overlaps between functions. All terms shown are significantly enriched (FDR-adjusted *p* < 0.05). **(B)** The 100 most highly expressed DEGs in N_BZ_ and N_GZ_ networks combined into a nonredundant set of 129 genes and their expression trajectory from EZ (center) to BZ (left) and GZ (right). Blocks of ribbons share a similar expression pattern. Colores represent the DEG group and expression strength. Functional processes enriched by these DEGs are listed with the number of genes contributed by each zone. **(C)** Specific genes among the top 100 highly expressed DEGs that showed contrasting contributions to the functions enriched in the N_BZ_ and N_GZ_ networks.

Building on this shared functional foundation, we next asked whether highly expressed genes could reveal processes that contribute to the morphological divergence between the two zones. We examined the most highly expressed genes as a representative subset of each network to assess its dominant transcriptional characteristics without evaluating every network component individually. Although cumulative changes across many genes likely contribute to the distinct functional profiles of the two zones, we reasoned that highly expressed genes would capture prominent components of the transcriptional programs required to withstand high salinity. These genes could be shared between zones, but we hypothesized that zone-specific phenotypes could arise from localized differences in the regulation of shared genes. To test this, we selected the 100 most highly expressed DEGs from each N_BZ_ and N_GZ_ network and consolidated them into a nonredundant set of 129 genes. We then tracked their expression patterns across the EZ-to-BZ and EZ-to-GZ transitions (Fig. 5B, Table S8).

The differences in expression of these shared genes include reversals in expression direction, with genes switching between up- and down-regulation, changes in the magnitude of expression strength (i.e. relatively low, moderate, and high among the top 100 highly expressed genes), and transitions between differentially expressed, and non-differentially expressed states (Fig. 5B). Nearly half of the shared genes maintained stable expression classes (i.e. their DEG status), whereas others diverged into distinct expression states. Although a large subset remained differentially expressed in both the N_BZ_ and N_GZ_ networks, a substantial portion was selectively deployed as DEGs in either the BZ or GZ (Fig. 5B, Table S8). These highly expressed genes were enriched in stress-response and signaling pathways but revealed zone-specific priorities. General stress responses were shared between the two zones, whereas oxidative stress responses were preferentially represented in the bulged zone, indicating a greater reliance on ROS-scavenging mechanisms. The two zones also differed in their hormonal responses relative to the control condition: the bulged zone preferentially recruited JA-associated processes, whereas the gap zone showed greater representation of auxin-associated processes, suggesting distinct hormonal regulation of the two zone-specific phenotypes.

Specific genes that differed in DEG status between the bulged and gap zones further revealed contrasting expression patterns associated with cell wall modifications, water and osmotic stress responses, oxidative stress homeostasis, cellular damage repair, and general stress tolerance in the two zones (Fig. 5C, Table S8). Notably, only two genes in this set, *EXTENSIN 3* (*EXT3*) and *CELL WALL-PLASMA MEMBRANE LINKER PROTEIN* (*CWLP*), showed diametric expression responses relative to the elongation zone-control: both were upregulated in the bulged zone but downregulated in the gap zone. *EXT3* encodes a well-characterized cell wall structural glycoprotein, whereas *CWLP* encodes a less-characterized cell wall–plasma membrane linker protein ^20,21^. Their coordinated but opposing regulation is consistent with a potential role in cell wall and plasma membrane-associated responses to localized salt-induced cellular stress. Together, these recurring functional patterns indicate that hormone-mediated developmental regulation, salt stress signaling and transport, cell wall integrity, and ROS homeostasis constitute major functional modules that are differentially deployed by the zone-specific transcriptional networks underlying the distinct bulged and gap zone phenotypes.

### Spatially distinct hormone networks accompany developmental remodeling under salt stress

To determine how transcriptional regulation of hormone pathways and hormone-responsive processes contribute to the distinct responses of the bulged and gap zones under salt stress, we examined the differently regulated genes associated with hormone biosynthesis, signaling, transport, and downstream responses in the N_BZ_ and N_GZ_ networks (Figure 6, Table S9). We previously showed that JA, IAA, and ABA accumulated differentially in the bulged and gap zones three days after salt exposure, when the transcriptional responses were measured (Fig. 3D, E). We therefore asked whether this hormone accumulation was accompanied by transcriptional changes in genes associated with these hormones. At the broader transcript-population level, genes associated with JA, IAA, and ABA showed shifts in mean expression under salt stress (Fig. 6A, Table S9). JA- and ABA-associated transcripts had higher mean expression in both the N_BZ_ and N_GZ_ networks, whereas the mean expression of IAA-associated transcripts decreased specifically in the bulged zone. In contrast, genes associated with gibberellic acid (GA) and cytokinin (CK) showed no significant transcriptional changes in either network (Fig. 6A, Table S9).

**Figure 6.**
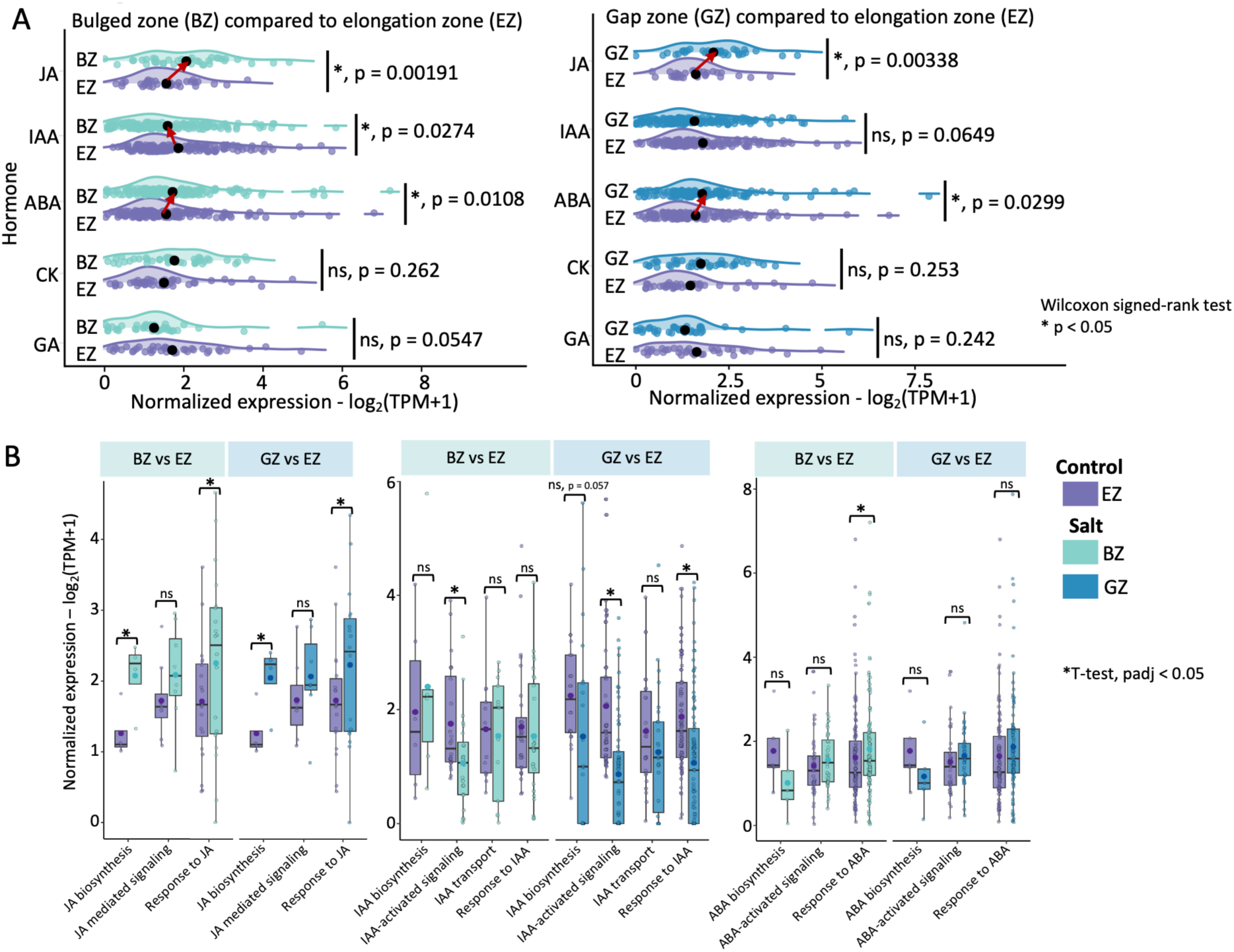
Spatial changes in hormone distribution are associated with distinct hormone-related processes in bulging and gap zones. **(A)** Distribution of expression levels for hormone-related differentially expressed genes (DEGs) between bulged and gap zones from the N_BZ_ and N_GZ_ networks. Black dots indicate mean expression values for each group, and colored dots represent individual DEG expression levels. Comparisons between root zones within each hormone group were performed using paired Wilcoxon signed-rank tests. Red arrows indicate the direction of transcriptional changes between control and salt-treated root zones for each hormone when the shift was significantly different. **(B)** Expression comparisons within each DEG group associated with hormone biosynthesis, signaling, transport, and response. The box plots are depicted with individual expression levels shown in small data points and mean expression in a large dot for each group. Statistical significance was assessed using paired t-tests followed by Benjamini–Hochberg correction for multiple testing. Boxes indicate the interquartile range with median lines. Significance was defined as adjusted *p* < 0.05 noted by *; *ns*, not significant. Boxes indicate the interquartile range with median lines. Hormones-jasmonic acid (JA), auxin (IAA), abscisic acid (ABA), gibberellic acid (GA), and cytokinin (CK). For all panels, gene sets were curated based on hormone-related Gene Ontology (GO) terms and filtered to include only DEGs identified in the salt-induced zone-specific networks N_BZ_ and N_GZ_ with TPM ≥ 1 in at least one root zone.

Examination of individual hormone-associated processes revealed notable differences between the zones (Fig. 6B, Table S9). JA biosynthesis genes were specifically induced in the N_BZ_ network, whereas genes associated with JA responses were significantly induced in the N_GZ_ network. For auxin, genes involved in auxin-activated signaling were suppressed in both zones relative to the N_EZ_ from the elongation zone, while genes known to respond to auxin were additionally suppressed in the gap zone. Interestingly, neither ABA biosynthesis nor ABA signaling showed significant transcriptional changes in the salt-induced zones by day 3. However, ABA-responsive genes were induced specifically in the bulged zone, suggesting that accumulated ABA contributes to the stress-responsive transcriptional network in this zone without requiring transcriptional induction of ABA biosynthesis or signaling. The transcriptional induction of JA biosynthesis in the bulged zone is consistent with our functional evidence that inhibition of JA biosynthesis with DIECA suppresses bulge formation under salt stress (Fig. 3A, B). Similarly, inhibition of root hair formation on the bulged zone by the auxin signaling inhibitor auxinole (Fig. S3) is consistent with the transcriptional pattern showing that auxin regulation is centered primarily on signaling rather than biosynthesis (Fig. 6B, Table S9). Together, these results indicate that although JA, IAA, and ABA accumulate in both salt-induced zones, transcriptional regulation of their biosynthesis, signaling, and downstream response pathways is spatially differentiated, contributing to distinct hormonal programs in the bulged and gap zones.

### Spatially distinct transcriptional networks balance cell wall expansion and structural integrity with ion, water, and energy homeostasis under salt stress

Both the bulged and gap zones must maintain cell wall integrity, although their distinct morphologies suggest different requirements: cells in the bulged zone expand, whereas those in the gap zone remain relatively uniform compared with the adjacent maturation zone and the preceding elongation zone (Fig. 2). We therefore examined all DEGs within the zone- and salt-specific N_BZ_ and N_GZ_ networks to identify genes associated with cell wall modifications. Only 10 DEGs met our search criteria (Figures 7A, B). Among these, *EXT3* was uniquely induced in the bulged zone, whereas *EXPANSIN 1 (EXPA1)* was uniquely induced in the gap zone; *EXTENSIN 21 (ETX21)* was induced in both zones relative to the elongation zone. *EXT3* is an essential cell wall structural protein required for physical reinforcement, whereas *EXPA1* in Arabidopsis is expressed in the root cap and lateral roots and its overexpression can arrest root growth ^21^. Their contrasting, zone-specific induction is consistent with coordinated regulation of cell wall properties to accommodate the distinct cellular states of the bulged and gap zones in *S. parvula*. The induction of *EXT21*, which remains poorly characterized in Arabidopsis, may further contribute to this balance between cell wall reinforcement and flexibility. We also noted that multiple *CELLULOSE SYNTHASE (CESA)* genes were downregulated in both the bulged and gap zones (Fig. 7A, B). The radial root swelling phenotype reported in *FEI* mutants under salt stress has impaired cellulose biosynthesis ^22^. However, the bulged zone phenotype we observed does not appear to result from reduced cellulose biosynthesis, as cellulose content increased in epidermal and cortical cells in the bulged and gap zones despite transcriptional downregulation of several *CESA* genes (Fig. 7B, C). In contrast, relative xylan abundance decreased in both zones (Fig. 7C).

**Figure 7.**
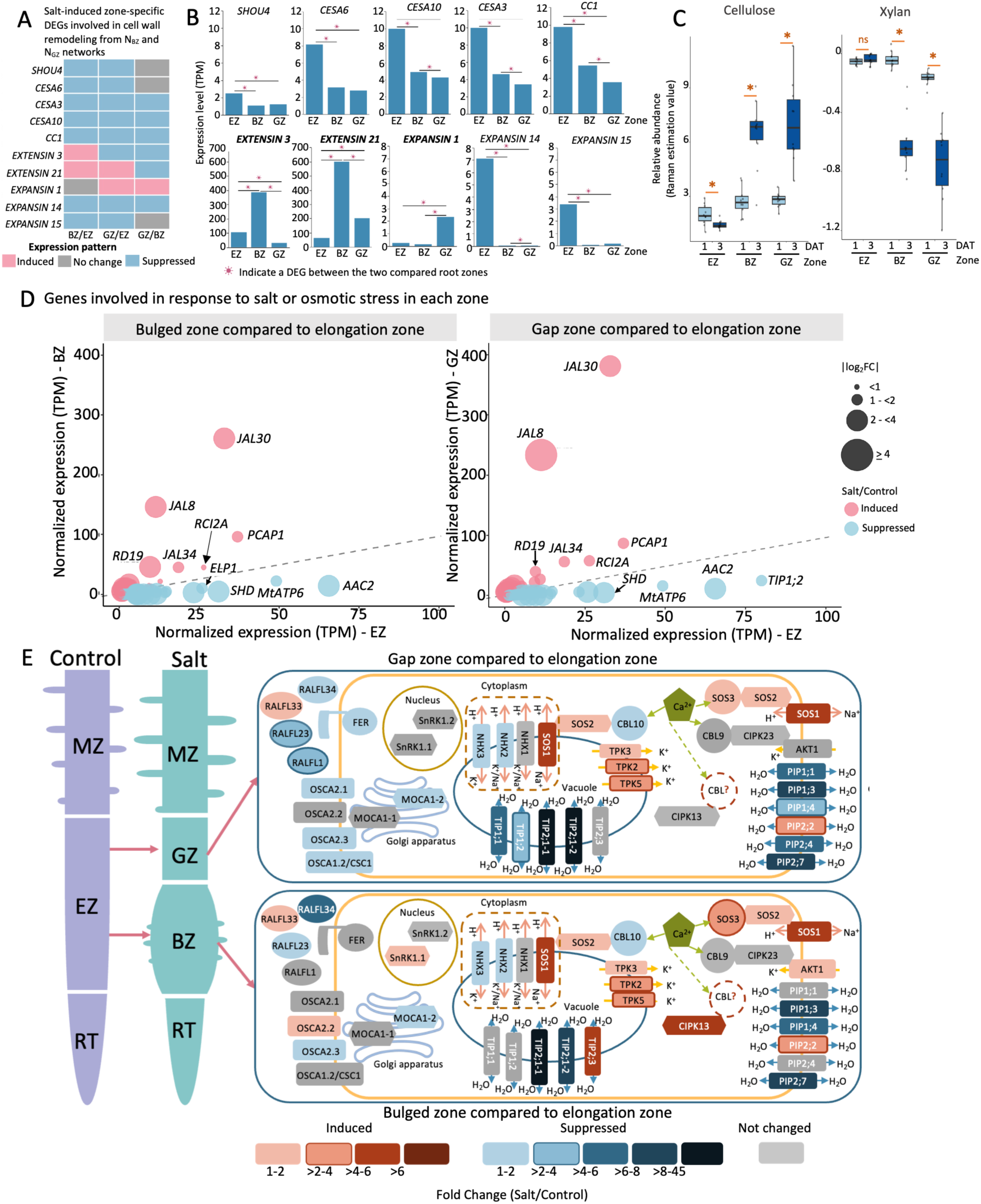
Spatial regulation of cell wall remodeling, ion and water balance is associated with distinct transcriptional strategies in the bulged and gap zones. (A) Differently expressed genes (DEGs) associated with cell wall remodeling in the N_BZ_ and N_GZ_ networks representing bulged (BZ) and gap (GZ) zones. Genes included in the heatmap were required to be differentially expressed in at least one zone compared to the elongation zone (EZ) and to have expression at TPM > 1 in at least one root zone. (B) Expression levels of selected cell wall– related genes across the three root zones (EZ, BZ, and GZ), highlighting zone-specific transcriptional responses under control and salt-treated conditions. Genes shown in this panel correspond to those in (A). (C) Cellulose and xylan content across root zones in the epidermal and cortical tissues using Raman spectroscopy at 1 and 3 days after 200 mM NaCl treatment (DAT). Each dot represents a data point. Measurements were obtained from at least three biological replicates, with a total of 10 data points per condition. Outliers were removed using Grubbs’ test (*p* < 0.05). Statistical comparisons between 1 and 3 DAT within each root zone, and among root zones within each time point, were performed using two-sample *t*-tests on individual measurements. *P*-values were adjusted using the Benjamini–Hochberg method. (D) The most highly expressed DEGs functionally annotated under response to salt and osmotic stress in the N_BZ_ and N_GZ_ networks. The top 10 genes with the highest TPM values and log₂FC ≥ 1 are labeled for each comparison. Bubble size represents |log₂FC|, and color indicates induced (pink) or suppressed (blue) expression under salt. (E) All DEGs captured in the N_BZ_ and N_GZ_ networks associated with salt-stress-related sensing, signaling, and transport mapped to their reported cellular localization. These include RALF–FER signaling, MOCA1-associated glycosyl inositol phosphorylceramide synthesis, OSCA-associated osmotic stress signaling, Ca²⁺ signaling, SOS pathway components, other Na⁺/K⁺/cation transport-related genes, and aquaporins. Colors indicate fold-change direction (up-brown and down-blue) and magnitude relative to the control expression level in EZ.

We next examined salt stress–responsive DEGs within the N_BZ_ and N_GZ_ networks that may support ion and water balance and thereby promote stress tolerance in the bulged and gap zones. Because maintaining these processes is energetically demanding, we also examined whether cellular processes that are less directly required for stress adaptation might be reciprocally reduced in these zones relative to the salt-unaffected elongation zone. We therefore focused on genes that were both highly expressed and strongly induced in each zone compared to the elongation zone (Fig. 7D). Several of the *JA-ASSOCIATED JACALIN-RELATED LECTINS (JALs*) were highly expressed and strongly induced in both the bulged and gap zones (Fig. 7D, Table S10). Among these, the most highly expressed *JAL30* and *JAL8* have been associated with defense and molecular chaperone functions, respectively ^23^. Overall, the transcriptional adjustments under high salinity suggest a coordinated balance between active stress tolerance and metabolic conservation. The concurrent induction of *PLASMA-MEMBRANE ASSOCIATED CATION-BINDING PROTEIN 1 (PCAP1), RESPONSIVE TO DEHYDRATION 19 (RD19)*, and *RARE-COLD-INDUCIBLE 2A (RCI2A)* represents a coordinated protective response, potentially involving calcium signaling at the plasma membrane, vacuolar protein turnover to recycle stress-damaged components, and membrane-potential stabilization to limit excess Na⁺ entry into the bulged and gap zones ^24–27^. In contrast, the simultaneous suppression of *ADP/ATP carrier 2 (AAC2), SHEPHERD (SHD)*, and *Mitochondrial F1F0-ATP synthase (MtATP6)* suggests reduced investment in mitochondrial energy transport, growth-associated protein functions, and ATP synthesis in these zones relative to the elongation zone ^28–30^ (Fig. 7D). Together, these changes indicate that the salt-induced zones prioritize protective and homeostatic functions while reducing localized energy-intensive processes associated with growth and metabolism.

To evaluate how these N_BZ_ and N_GZ_ networks in the root zones transcriptionally adapt to compartmentalize Na⁺, regulate water flux, and maintain potassium balance, we mapped all the differently expressed genes associated with salt sensing, osmosensing, water transport, Na⁺ transport, and K⁺ homeostasis across the bulged and gap zones (Fig. 7E). Under salinity stress, both zones activated the core Salt Overly Sensitive (SOS) pathway ^31^ to restrict cytosolic Na⁺ accumulation. Concurrently, both zones induced calcium-activated, *K⁺-SELECTIVE TWO-PORE POTASSIUM* (*TPK/KCO*) channels ^32,33^ on the tonoplast to potentially preserve cytoplasmic K⁺ homeostasis. However, the transcriptional processes facilitating K⁺ uptake and osmotic stress adjustments diverged between the two zones. The bulged zone exhibited an induction of the potassium channel *AKT1* and its potential regulator, *CBL-INTERACTING PROTEIN KINASE 13 (CIPK13)* (Fig. 7E). *CIPK13* is highly expressed during root hair morphogenesis in Arabidopsis ^34,35^. Its upregulation aligns with the initiation of root hairs observed in the bulged zone (Fig. 2A) as well as the need to increase K⁺ uptake under salt stress ^1^. Furthermore, the bulged zone uniquely induced the hypo-osmosensitive calcium-permeable channel *REDUCED HYPEROSMOLALITY-INDUCED CA2+ INCREASE 2.2 (OSCA2.2)* ^36^, indicating that these cells experience local hypo-osmotic stress (Fig. 7E). Synergistically, the tonoplast-localized aquaporin *TIP2;3* was highly and specifically induced in the bulged zone, whereas other water channels remained suppressed or at baseline levels across both zones (Fig. 7E). The energy-sensing kinase *SnRK1*, which acts as a master regulator of primary metabolism ^37^, was also induced exclusively in the bulged zone, likely acting to arrest local growth-related energy expenditure and activate downstream ABA-responsive networks. This aligns with the tissue-specific accumulation of ABA, which was higher in the epidermis and cortex cell layers of the bulged zone at day 3 compared to the elongation zone at control levels (Fig. 3E). Together, these results demonstrate that while both zones successfully achieve salinity tolerance and coordinately contribute to the recovery of primary root growth, they do so through distinct localized stress-sensing mechanisms and divergent transcriptomic responses.

### Coordinated metabolic and transcriptional responses limit ROS accumulation and cell death under high salinity

One of the most distinguishing features when transitioning from the elongation zone to a salt-adapted state in the bulged and gap zone is the control of cell death (Fig. 1C). A primary contributor to cell death would be osmotic stress and excessive ROS generation expected under salt stress. Therefore, we next examined whether there was an oxidative burst reflected by excessive build up ROS and if not, whether a boost in antioxidants and osmoprotectants could explain how the cells prevented excessive cell death at 200 mM NaCl when the primary root is actively growing (Fig. 1B). Although localized superoxide production is a known developmental signal that initiates processes such as lateral root formation, our aim was to distinguish these spatially restricted ROS signals from excessive and broadly distributed ROS accumulation indicative of oxidative stress and salt susceptibility. We used H₂DCFDA (2′,7′-dichlorodihydrofluorescein diacetate) which enters cells readily, as a fluorescent probe to assess the general intracellular oxidative activity/ROS levels ^38^. Roots treated with high salt showed ROS levels comparable to control roots, whereas roots treated with extreme salinity showed strong ROS accumulation across the root axis (Figures 8A, B). These results suggest that *S. parvula* limits excessive ROS accumulation under high salinity, which may help protect root cells from oxidative damage.

**Figure 8.**
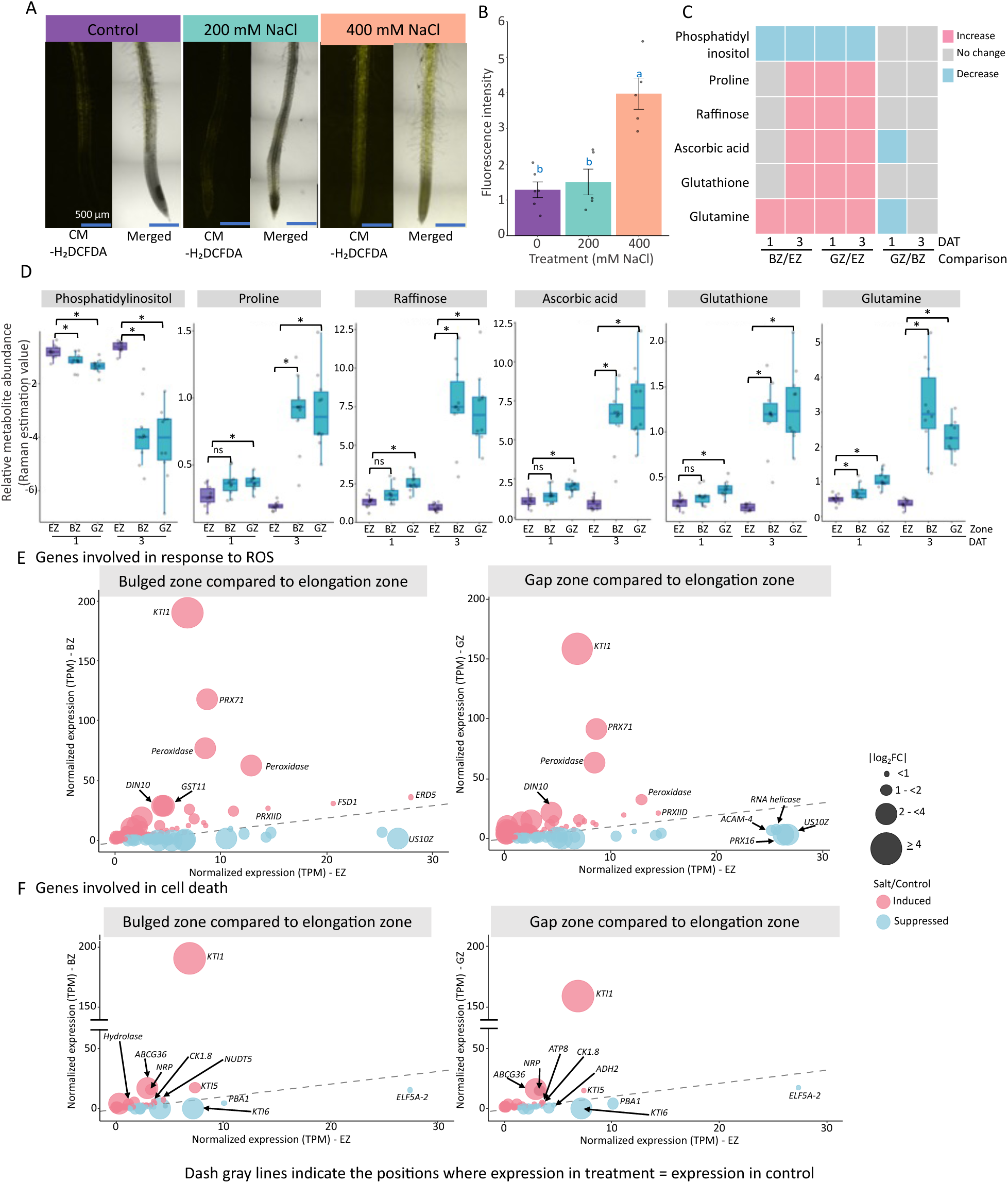
Coordinated metabolic and transcriptional responses limit ROS accumulation and cell death under high salinity. **(A)** *S. parvula* roots treated with control solution, 200 mM NaCl, or 400 mM NaCl for 3 days after treatment (DAT), stained with H₂DCFDA (2′,7′-dichlorodihydrofluorescein diacetate) to detect general ROS accumulation. Confocal images were digitally stitched from overlapping frames to capture the extended vertical length of the primary root. **(B)** Quantification of ROS signal intensity along the root axis. **(C)** Differential accumulation of osmolytes and antioxidants measured by Raman spectroscopy at 1 DAT and 3 DAT in epidermal and cortical cell layers of the control elongation zone (EZ), salt induced bulged zone (BZ), and gap zone (GZ). **(D)** Relative abundance of individual metabolites across spatial and temporal progression. Statistical comparisons were performed using Welch’s two sample t test, with p values adjusted using Holm’s correction (ns-not significant, *-*p_adj_* < 0.05). **(E)** Normalized expression of DEGs associated with ROS responses in the BZ and GZ from the N_BZ_ and N_GZ_ networks, compared with the control EZ. **(F)** Normalized expression of DEGs associated with cell death in the N_BZ_ and N_GZ_ networks, compared with the control EZ. Bubble size represents |log₂FC|, and color indicates genes induced (pink) or suppressed (blue) by salt treatment. The top 10 genes with the highest TPM values and log₂FC ≥ 1 are labeled for each comparison.

To identify metabolic changes associated with this low-ROS state under high salinity, we quantified metabolites with established protective roles as osmoprotectants and antioxidants (Fig. 8C, D). We also quantified phosphatidylinositol as a molecular indicator of cellular stress, because reduced phosphatidylinositol levels have been associated with activation of the SOS pathway and Ca²⁺ signaling in Arabidopsis roots under salt stress ^39^. We observed a clear reduction in phosphatidylinositol levels in both the bulged and gap zones relative to the elongation zone, where phosphatidylinositol remained at high basal levels (Fig. 8D). Consistent with the low-ROS state observed under high salinity, several metabolites with established roles in osmotic protection, antioxidant defense, and metabolic reallocation were significantly increased in both the bulged and gap zones relative to the elongation zone (Fig. 8C, D). The observed increased proline may lower cytosolic water potential to maintain turgor and water uptake, while also stabilizing membranes and directly scavenging reactive oxygen species ^40^. The increased raffinose may further protect cellular structures by promoting vitrification and limiting physical damage during osmotic stress ^41^. The coordinated accumulation of ascorbic acid and glutathione provides enhanced capacity for ROS detoxification through the ascorbate– glutathione cycle, with ascorbate serving as a major antioxidant and glutathione supporting its regeneration and continued activity ^42^. Glutamine, a major nitrogen donor for the synthesis of stress-protective metabolites such as proline ^43^, was also elevated, potentially supporting the metabolic shift from growth toward stress tolerance (Fig. 8D). Together, these metabolic changes indicate that the bulged and gap zones reinforce osmotic protection, antioxidant capacity, and resource allocation to maintain cellular integrity under high salinity. We next examined salt stress–responsive genes concordantly induced within the N_BZ_ and N_GZ_ networks associated with ROS detoxification and prevention of cell-death regulation in the bulged and gap zones. In total, 261 DEGs in the bulged and 239 DEGs in the gap zones were associated with ROS responses, while 89 DEGs in each zone were associated with cell death, with partial overlap between the two zones. Notably, *GLUTATHIONE S-TRANSFERASE 11 (GST1)* and *FE SUPEROXIDE DISMUTASE 1 (FSD1)* were specifically induced in the bulged zone (Fig. 8E, Table S10). *GST11*, a glutathione S-transferase, can contribute to ROS detoxification by conjugating reduced glutathione to cytotoxic lipid peroxides under salt stress ^44^. *FSD1*, an iron-containing superoxide dismutase, converts superoxide radicals to H₂O₂ and O₂ to balance localized ROS levels and preserve root hair tip growth under high salinity ^45^.

We next examined the transcript levels of highly expressed genes involved in modulating cell survival and death. Among these, *KTI1*, a Kunitz trypsin inhibitor previously shown to limiting programmed cell death (PCD) progression and lesion formation ^46^, was strongly induced in both the bulged and gap zones, with expression levels exceeding 20-fold relative to the control elongation zone. Concurrently, a robust transcriptional adjustment of ROS metabolism was observed in these salt-induced zones, suggested by the differential expression of several peroxidase-related genes, including *PRX71, PRXIID, PRX16*, and multiple uncharacterized *peroxidases* (Fig. 8E, F, Table S10). In contrast, *RIBOSOMAL PROTEIN US10Z (uS10z)*, which encodes a small ribosomal subunit protein, and *EUKARYOTIC ELONGATION FACTOR 5A-2 (ELF5A-2/eIF5A-2)*, a known positive regulator of PCD ^47^, were both downregulated in the bulged and gap zones. Collectively, these coordinated expression profiles indicate a localized prevention of excessive cell death under high salinity, driven by the concurrent activation of antioxidant networks alongside the targeted suppression of pro-PCD cellular machinery.

## DISCUSSION

Increasing salt concentrations in the growth medium cause either irreversible root damage or growth adjustments in all plants ^48,49^. Salt-adapted plants withstand higher salt concentrations and maintain growth and cellular integrity compared with salt-sensitive plants ^5,50^. Previous studies identified vulnerable points that can lead to cell death and growth inhibition in eudicot root systems, using Arabidopsis as a model ^51–53^. Using the salt-adapted model *Schrenkiella parvula*, our study examined how roots withstand increasing salinity by reprogramming their developmental trajectories to sustain primary root growth under high salt stress. Our findings demonstrate that root growth can recover at 200 mM NaCl, revealing a capacity to sustain growth under salinity levels generally considered severely inhibitory to crops.

Root responses to salt occur across multiple timescales, ranging from seconds to weeks ^53,54^. We focused on the stable, stress-adapted state in which root growth had resumed following the transient growth pause in *S. parvula* (Fig. 1). This timing provides an opportunity to identify cellular and developmental mechanisms that may be absent or obscured in the initial stress-response phase. Studies in Arabidopsis have shown that high salinity inhibits root tip growth by halting cell expansion, driving ROS accumulation, and inducing excessive cell death, particularly within the elongation zone ^51,52,55^. In contrast, *S. parvula* sustains growth recovery through a distinct developmental transition within its elongation zone, characterized by a "bulged zone" and a "gap zone" (Fig. 2). The newly established developmental zones exhibited extensive transcriptional responses to salt treatment, as indicated by their large numbers of DEGs. By systematically dissecting the zone-specific transcriptomes, we resolved general stress responses and pre-existing developmental responses from responses uniquely activated by salt within these localized zones (Fig. 4E-G, 5, Tables S5, S6). These specialized zones exhibit localized differential accumulation of JA, ABA, and auxin that correspond with their spatially distinct transcriptional responses (Fig. 3D, E, 6). These zones also undergo changes consistent with developing structurally resilient cell walls and membranes that resist salt-induced collapse while maintaining regulated water transport, in agreement with the corresponding transcriptional signatures (Fig. 7). Concurrently, ROS accumulation remains suppressed at levels comparable to non-saline controls, accompanied by transcriptional induction of antioxidant pathways and increased accumulation of compatible osmolytes and antioxidants (Fig. 8). Collectively, we present a spatiotemporally coordinated program of cellular adjustments that actively shields primary roots from known cellular vulnerabilities, optimizing salinity tolerance.

Elevated levels of ABA, JA, and auxin are widely documented to suppress cell division and expansion, thereby inhibiting overall root growth ^56–59^. Under salinity stress, in salt-sensitive plants, increased ABA acts through the receptor-like kinase FERONIA (FER) or downstream auxin pathways to arrest root cell elongation ^60,61^. Similarly, osmotic stress induces JA-responsive *JASMONATE ZIM-DOMAIN (JAZ)* genes, initiating signaling cascades that restrict cell elongation ^57^. Notably, these growth-inhibitory pathways fail to explain how *S. parvula* sustains continuous growth despite high accumulation of these hormones in the bulged and gap zones (Fig. 3D, E). Instead, our findings reveal three major deviations from standard salt-sensitive responses. First, cell numbers remain stable between the elongation zone and the salt-induced bulged and gap zones (Fig. 2C). We observed neither the induction of *FER* nor the ABA-mediated transcriptional repression of B-type *ARR2*, a transcription factor in cytokinin-signaling required for cell elongation (Fig. 7E). Additionally, cytokinin-associated transcriptional pathways remained largely unchanged in *S. parvula* (Fig. 6A). Second, the osmotic stress- and JA-responsive *JAZ (Jasmonate ZIM-domain)* genes were not induced, which aligned with our finding that osmotic stress alone was insufficient to trigger this developmental transition in *S. parvula* (Fig. 2D). Third, JA accumulation appears to mediate this developmental reorganization through a highly cell-layer- and zone-specific manner (Fig. 3D, E). Members of the *JACALIN-RELATED LECTIN* (*JAL*) family stand out among the most highly upregulated genes in the bulged and gap zones (Fig. 7D). While their roles under salinity remain poorly characterized, the Arabidopsis orthologs of *S. parvula JAL30* (also known as *Jasmonic Acid-Induced Protein/JIP*) and *JAL8* are associated with cellular defense responses ^62,63^. Furthermore, while JA is reported to impair salinity tolerance in salt-sensitive plants by repressing *CATALASE 2 (CAT2)* expression ^64^, we observed the opposite behavior: *CAT2* is among the most highly induced genes specifically restricted to the gap zone, facilitating localized ROS suppression (Fig. 5C). Notably, genes involved in JA biosynthesis were highly expressed in the bulged zone (Fig. 6B), suggesting that the bulged zone may be a site of local JA production and supports a model in which JA contributes to the formation of the bulged zone under high salt. Exogenous MeJA induced root bulging, whereas inhibition of JA biosynthesis under salt stress suppressed bulging (Fig. 3A, B). JA is known to act through cell-type-dependent programs where JA signaling is shaped by a complex network of negative regulators within Arabidopsis ^65^. Our results highlight even a more diverged regulatory capacity for JA in a salt-adapted plant that converts a canonical growth inhibitor into a mediator of salinity tolerance facilitating continuous growth under high salt.

In Arabidopsis, inadequate response to salt can trigger salt-induced root swelling, while exogenous ABA application can mitigate the cellular damage caused by uncontrollable cell expansion and swelling ^66^. Additionally, temporal mapping of transcriptional responses to salt indicates that ABA-related responses are broadly activated across root cell layers during the early quiescent phase of salt exposure, whereas JA-related pathways become prominent during the later recovery phase, spanning most cell layers with the exception of the epidermis ^53^. In contrast, ABA appears to have a divergent regulatory role in *S. parvula*, which aligns with a previous study examining cell-type-specific responses between Arabidopsis and *S. parvula* under salt- and ABA-treated conditions ^14^. While our results support a model in which ABA can act as a broad mediator of salinity and osmotic stress responses to regulate systemic water and ion homeostasis (Table S10), ABA may also contribute to the signaling cascade that coordinates zone-specific responses, including osmotic stress regulation and Na⁺ transport through the SOS pathway (Fig. 7E).

Contrary to the general expectation that salinity causes hyperosmotic stress leading to intracellular water loss and a high vulnerability to plasmolysis, the *S. parvula* bulged zone appears to experience a localized hypoosmotic environment, as suggested by the significant induction of the hypoosmotic sensor *OSCA2.2* (Fig. 7E) ^36^. Given that cells under hypoosmotic or high turgor pressure expand laterally, tight regulation of water transport is essential to prevent excessive hydrostatic pressure from fracturing the cell wall. Consistent with this requirement, multiple plasma membrane- and tonoplast-localized aquaporins are notably downregulated within the bulged zone (Fig. 7E). This targeted downregulation indicates that the bulged state in *S. parvula* is different from the generalized root swelling observed in salt-sensitive plants ^52,67^.

Concurrently, the bulged zone accumulates high levels of Na^+^ (Fig. 1C). The observed reciprocal induction of the SOS pathway, which removes Na⁺ from the cytosol, and K⁺ transporters, which promote K⁺ uptake into the cytosol, may counteract cytosolic ion toxicity and maintain a favorable K⁺/Na⁺ ratio in the bulged zone (Fig. 7E). Our results suggest that *S. parvula* maintains a localized zone of high-water availability while regulating cell expansion to prevent uncontrolled swelling and initiating root hair development within the bulged region, as indicated by its hypoosmotic response (Fig. 7E). This microenvironment may establish bidirectional spatial cues, with the region immediately above the bulged zone promoting lateral root initiation and growth, while the uncompromised tissues below sustain cell division and primary root tip growth. Together, these coordinated growth responses may allow *S. parvula* to maintain both primary root elongation and lateral root development, thereby enhancing growth resilience under high salt stress at 200 mM NaCl (Fig. 1B, 2E). Notably, whereas wild-type *Arabidopsis* does not exhibit this adaptation under high salinity, the cell-wall mutant *KORRIGAN1 (KOR1)* accumulates elevated JA in seedling roots, where enlarged cortex cells compress inner tissues and induce JA biosynthesis ^68^. Mechanical stress is known to activate JA signaling ^69^ and JA together with auxin, regulates lateral root positioning and initiation ^70,71^. Consistent with this mechanism, JA and auxin remained elevated in the epidermal and cortical layers of the *S. parvula* gap zone 3 days after salt treatment (Fig. 3E), coinciding with localized lateral root emergence (Fig. 2E). Moreover, lateral roots consistently emerged immediately above the bulged zone, suggesting that the spatially restricted cellular remodeling may provide a positional cue for lateral root initiation.

Equally important to sustain undamaged roots is the maintenance of cell wall integrity, particularly in the bulged zone, where cells are under high turgor pressure. Our findings suggest that *S. parvula* achieves this through localized, zone-specific transcriptional regulation of cell wall-modifying genes including *EXT3* and *EXT21* that may provide critical structural reinforcement ^20,72–75^ in addition to the increased deposition of cellulose under high salinity (Fig. 7A-C). Additionally, the gap zone uniquely induces *EXPA1* (Fig. 7A, B). In *Arabidopsis*, overexpression of *EXPA1* causes root growth arrest ^21^. Therefore, the spatially distinct induction of *EXT3, EXT21,* and *EXPA1* suggests that the two salt-induced zones differentially remodel cell wall properties rather than simply experiencing salt-induced wall damage. This spatial reorganization contrasts with the cell wall defects reported in Arabidopsis under salt stress ^22^.

A characteristic feature of *S. parvula* roots is their remarkable capacity to suppress ROS accumulation and limit cell death (Fig. 1C, 8A, B). Unlike salt-sensitive plants, *S. parvula* maintains intracellular ROS levels close to those of non-saline controls under high salinity (Fig. 8A, B). This protection is supported by coordinated transcriptional and metabolic adjustments involving compatible osmolytes, antioxidants, and suppressors of programmed cell death (Fig. 8C–F). By preventing excessive ROS accumulation, *S. parvula* may preserve ROS signaling required for developmental processes that promote continued root growth, including lateral root emergence in the gap zone (Fig. 2E). For example, the *Arabidopsis* ortholog of *LATERAL ORGAN BOUNDARIES DOMAIN 11 (LBD11)*, which is associated with ROS-mediated regulation of meristematic activity ^51^, is induced in the gap zone of *S. parvula*, whereas *LBD11* is downregulated by salt stress in *Arabidopsis* roots, where its suppression is associated with root growth arrest. Our analysis identified many genes that were preferentially induced in specific salt-adapted zones (Table S5), providing candidates for uncovering regulatory mechanisms that may distinguish salt-adapted from salt-sensitive plants. Together, these findings indicate that the bulged and gap zones function as metabolic and transcriptional hubs that integrate ROS detoxification, osmotic protection, and cell-survival pathways while preserving developmental signaling needed to sustain primary root growth under high salinity.

Functional studies will be needed to define the specific transcriptional regulatory cascades from upstream transcription factors to their downstream targets that underlie these responses in *S. parvula*. Importantly, this study provides the spatiotemporal context needed to move beyond individual candidate genes by defining when and where these genes are expressed and the transcriptional partners with which they are coordinated within salt-induced developmental networks. Resolving these unique regulatory configurations provides a molecular roadmap for identifying tissue-specific strategies to circumvent cellular vulnerabilities and ultimately engineer improved salinity tolerance in crops.

## MATERIALS AND METHODS

### Plant growth conditions for physiological assessments and RNA-seq

*Schrenkiella parvula* (ecotype Lake Tuz) seeds were sterilized and stratified at 4°C as previously described ^76^. These were placed on quarter-strength Murashige and Skoog medium solidified with 0.8% (w/v) phytoagar (¼ MS plates) for germination. Seven-day-old seedlings were transferred to ¼ MS plates supplemented with 0 to 400 mM NaCl and grown for up to seven days, depending on the experiment. Plates were kept at 22°C to 24°C in a growth chamber with a 16-h-light/8-h-dark cycle; 100-150 µmol m^-2^ s^-1^ light intensity.

### CoroNa Green and Propidium Iodide staining

To examine Na⁺ accumulation and cell death in roots, 7-day-old *S. parvula* seedlings were transferred to ¼ MS plates supplemented with 0, 200, or 400 mM NaCl and treated for 3 days. At each time point, roots were excised for staining. To visualize Na⁺ accumulation, roots were stained with 25 µM CoroNa™ Green (Invitrogen, CAS: C36675) for 3 h, as described by ^77^. To distinguish dead cells from live root cells, roots were stained with 10 µg/mL propidium iodide (PI; Invitrogen, CAS: P1304MP) for 30s. After staining with CoroNa Green and/or PI, roots were rinsed with distilled water to remove excess dye, mounted in water, and visualized by confocal microscopy. Cells in which PI had penetrated into the cytosol were considered dead.

### Measurement of cortical cells

10-day-old seedlings, which had been treated with 0 or 200 mM NaCl for 3 days, were prepared and stained with 0.1% Calcofluor White (Sigma Aldrich, CAS: 18909) as described in ^78^. Samples were imaged on Leica SP8 Confocal Microscope. To compare the cortical cell size along the root axis between control and 200 mM NaCl treatments, lengths and widths were measured from the maturation zone to the onset of elongation (i.e., length is more than width) for control, and bulge zone for salt-treated seedlings. Further, to determine if the cortical cell number changes between control and salt-treated seedlings, we counted the total number of cortical cells from the maturation zone to the meristem.

### Detection of reactive oxygen species (ROS)

For the detection of superoxide, 7-day-old *S. parvula* seedlings were treated with 200 mM NaCl, and roots were stained with Nitroblue Tetrazolium Chloride (NBT) (Sigma-Aldrich, CAS: 298-83-9) after 1 and 3 days of treatment as described in ^56^. In short, roots of treated seedlings were incubated in 20 mM phosphate buffer (pH 7.4) containing 0.05% (mass/volume) NBT in 1.5 mL tubes for 1 h at 37°C in the dark. After incubation, the staining solution was removed, and roots were immersed in 70% ethanol for 30 minutes before being imaged. To visualize general ROS accumulation in roots, 7-day-old *S. parvula* seedlings were transferred to ¼ MS plates supplemented with 0, 200, or 400 mM NaCl and treated for 3 days ^79^. Roots were excised and stained with 20 µM of 5-(and-6)-chloromethyl-2’,7’-dichlorodihydrofluorescein diacetate, acetyl ester (CM-H₂DCFDA) (Invitrogen, CAS: C6827) in the corresponding ¼ MS medium with the appropriate NaCl concentration for 30 min at room temperature in the dark. After staining, roots were rinsed with distilled water to remove excess dye, mounted in water, and visualized by confocal microscopy. Fluorescence signal intensity was quantified along the root axis to compare ROS accumulation across treatments.

### Osmotic and ionic stress treatments

7-day-old *S. parvula* seedlings were plated in ¼ MS media supplemented with 400 mM Mannitol (VWR, CAS: 69-65-8) to induce osmotic stress. *S. parvula* seedlings were treated with a series of salt 200 mM KCl or NaCl to test for ionic stress. Roots were cut, washed, and then mounted with distilled water before being imaged.

### Phytohormone treatments

7-day-old *S. parvula* seedlings were plated in ¼ MS media with or without added 200 mM NaCl and supplemented with different phytohormones or their inhibitors in different concentrations as listed: 50 μM Methyl jasmonic acid (MeJA, Sigma, CAS 39924-52-2), 1mM Sodium diethyldithiocarbamate (DIECA – JA biosynthesis inhibitor, ThermoFisher, CAS 20624-25-3); 0.1 μM Auxin (IAA, Sigma, CAS 87-51-4); 5 μM Auxinole (auxin signaling inhibitor, MedChemExpress, Cat. No. HY-111444). For time-lapse video, 10 μM ABA (Sigma-Aldrich, CAS 14375-45-2) was applied.

### Root imaging and image processing

Fluorescence imaging was performed using a Leica TCS SP8 confocal microscope at the LSU Shared Instrumentation Facility. CoroNa Green was excited with the 488 nm laser, and emission was detected between 502 and 516 nm. Propidium iodide (PI) was excited with the 561 nm diode laser, and emission was detected between 580 and 680 nm. CM-H₂DCFDA was excited at 492-495 nm, and emission was detected between 517 and 527 nm. The wavelengths for Calcofluor White excitation and emission were 405 nm and 425-475 nm, respectively. All confocal images were captured at 12-bit depth using an HC PL APO CS2 40X/1.3 oil objective, a laser power setting of 1.0, and a pinhole 1.2.

A VWR Professional Plus Phase Trinocular Microscope (Cat. No. 76122-406) was used to image roots stained with NBT and roots from osmotic, ionic, and phytohormone treatment assays.

For the primary root growth assessments, plant growth plates were scanned using the Epson Perfection V600 Photo scanner at 1200 dpi. Primary root growth was calculated as the change in root length between time points.

For time-lapse imaging, seedlings were grown as described above and used at 5–7 days post-germination. Seedlings were transferred onto agar pads prepared with the same composition as the growth medium. Using a razor blade and forceps, each agar pad was inverted and gently placed onto a microscope chamber slide (Lab-Tek II, Chambered, cat no. 155360), such that the resulting assembly consisted of the slide, the plant, and the agar pad in that order (slide–plant– agar). The chamber was then sealed with 3M Millipore tape and mounted on an inverted microscope (Leica, DMi8). Brightfield images were acquired every 15 min over a total imaging period of 16h at 20X magnification.

Image processing and quantification were performed using Fiji/ImageJ2 version 2.9.0/1.53t. Fiji was used to measure primary root length from scanned plate images, cortical cell morphology, and quantify fluorescence intensity.

### Tissue sampling for transcriptome and targeted metabolite quantification

To collect samples for RNA-seq or quantification of hormones using LC-MS, 7-day-old *S. parvula* seedlings were transferred to ¼ MS agar plates supplemented with either 0 or 200 mM NaCl and maintained in the growth chamber under the conditions described above for 1 and 3 days of treatment. All samples were harvested and sectioned at the same time window to minimize circadian effects.

Root tissue zones were sectioned and stored as described by Brady et al., 2017 ^80^ for RNA-seq, with minor modifications. For hormone quantification, sectioned tissues were stored in deionized water. After removal from treatment plates, roots were mounted in ¼ MS liquid medium supplemented with the corresponding NaCl concentration. Root zones were dissected using double-edged razor blades, and dissected root sections were transferred with fine needles into tubes containing RNA extraction buffer/deionized water kept on ice. Samples were immediately frozen after harvested and stored at -80°C until RNA/hormone extraction. Under control conditions, roots were dissected into three zones: root tip (RT), elongation zone (EZ), and maturation zone (MZ). Under salt treatment, roots were dissected into four zones: root tip (RT), bulging zone (BZ), gap zone (GZ), and maturation zone (MZ). Root-zone boundaries were defined based on cortical cell morphology and position along the root axis. The RT was collected from the root apex to the point where cortical cells first showed elongation along the vertical/longitudinal axis. The EZ was defined as the region above the RT where cortical cells were visibly elongated, typically reaching approximately two-fold greater longitudinal length than cells in the basal RT region ^81,82^. The MZ was collected from the first root hair-bearing cell and extended up to 1 cm above this point, and this section did not include lateral roots ^83^. For salt-treated roots, the BZ was collected from the beginning of visible cortical cell expansion to the transition with the GZ. The GZ was defined as the root-hairless region containing narrow cells between the BZ and the first root hair-bearing cell marking the beginning of the MZ. Approximately 50-100 root sections were collected per sample, with 3–4 biological replicates per condition.

### RNA-seq library preparation, sequencing and initial data processing

Total RNA was extracted using the RNeasy Plant Mini kit (Qiagen, Hilden, Germany), followed by an additional DNase treatment to remove genomic DNA contamination. Due to the small sample size and low RNA concentrations obtained from individual root sections, RNA integrity and quantity were assessed using an Agilent 2100 Bioanalyzer (Agilent Technologies, Santa Clara, CA, USA) at the LSU Genomics Facility prior to library preparation. RNA Integrity Number (RIN) values ranged from 6.2 to 9.6, with a mean RIN of 8.27, indicating that the majority of samples were of high quality and suitable for downstream RNA-seq analyses. 3-4 biological replicates per condition were used for RNA-seq libraries.

RNA-seq libraries were prepared at the Roy J. Carver Biotechnology Center, University of Illinois at Urbana Champaign. Libraries were barcoded and sequenced on one lane of NovaSeq X Plus, generating 11-16 million high-quality 150-nucleotide paired-end RNA-seq reads per sample.

Raw paired-end reads were quality checked, adapter-trimmed, and filtered using fastp ^84^, then mapped to *S. parvula* genome v4.1 using STAR aligner ^85^. Uniquely mapped reads were counted using featureCounts ^86^, and differentially expressed genes (DEGs) were identified using DESeq2 ^87^ with a FDR-adjusted p-value cutoff of 0.05. Read counts were then normalized to TPM for downstream visualization and comparison of relative transcript abundance when relevant.

To identify enriched functional categories in the RNA-seq datasets, Gene Ontology (GO) enrichment analysis was performed using BiNGO ^88^ implemented in Cytoscape (Version 3.10.3). GOMCL was subsequently used to cluster related enriched GO terms and generate a non-redundant summary of functional categories and the genes associated with each enriched group^89^.

### Dissection of total transcriptomes into T, H, U, and N modules

To identify salt-induced zone-specific responses, regular developmental differences among root zones were separated from salt-responsive transcriptional changes (Fig. 4E, S4A). Developmental DEG sets were first defined for each control root zone by comparing each target zone to the combined expression profiles of the other two control zones, identifying 11,266 RT-dependent DEGs, 4,865 EZ-dependent DEGs, and 11,918 MZ-dependent DEGs (Fig. S4C). The whole-root salt-responsive genes, defined as set U, was selected by DEGs shared across salt-versus-control comparisons and regulated in the same direction (Figs. S4B and S4D). Because the BZ and GZ are both derived from the control EZ, genes that differed significantly between BZ and GZ were excluded from set U. Thus, set U, which had 1,387 DEGs, represents broad salt-responsive genes shared across the entire root rather than BZ- or GZ-specific responses. For each target root zone, salt-dependent and zone-dependent DEG sets (set H) were then defined by combining up- and down-regulated DEGs from the corresponding salt versus control comparison and removing set U (Fig S4E). Finally, salt-induced zone-specific DEG sets, defined as set N, were obtained by subtracting the corresponding developmental DEG set (set T) from these salt-dependent, zone-dependent DEG sets (set H) (Fig. 4E, S4F). Therefore, set N represents genes specifically associated with salt responses in each root zone after excluding both normal developmental differences and whole-root salt responses.

### Construction of T, H, U, and N gene networks

The lists of differentially expressed genes (DEGs) and their corresponding normalized transcript abundances (transcripts per million; TPM) from the T, H, U, and N modules within each zone were used to calculate pairwise Pearson correlation coefficients. Only gene pairs showing strong correlations (|r| ≥ 0.90) with p ≤ 0.001 were retained for Louvain community detection and co-expression network analysis. For network visualization, all DEGs assigned to the same Louvain community were collapsed into a single node, with node size representing the number of genes within that community. Communities belonging to the same connected metacommunity (i.e., a higher-level network grouping) were linked by edges to visually represent their shared membership in a metacommunity. Communities that did not form a metacommunity were kept as single nodes. The DEGs that did not meet the cutoff to be considered co-expressed were represented by the smallest nodes without edges (i.e. did not form into co-expressed gene communities).

Our goal was to identify processes that are prioritized within coordinated yet spatially restricted programs, which may be obscured when each zone is analyzed independently. The T, H, U, and N networks for each zone were summarized by the number of DEGs belonging to (i) multi-gene metacommunities, defined as connected components containing more than one Louvain community; (ii) multi-gene single-communities, defined as components containing only one Louvain community but with multiple genes; and (iii) single-gene nodes, representing individual genes with low correlated expression to all other genes in the network (Table S6). The resulting community-level node and edge networks were visualized using Cytoscape v3.10.4.

### Identification of orthologs between *S. parvula* and *A. thaliana*

Genome annotations for *S. parvula* version 4.1 and *A. thaliana* genome version 11 (https://www.araport.org/) were used for ortholog identification. Orthologous gene pairs as best reciprocal hits between these two species were identified using the CLfinder-OrthNet pipeline with default settings ^90^. To account for lineage-specific gene duplications in both species, orthologous gene pairs were searched reciprocally between the two-species using BlastP with an e-value of 1e-5 and MMseqs2 ^91^ with an equivalent e-value cutoff. These pairs were further filtered using OrthoFinder ^92^ with granularity -I of 1.6 and were added back to the CLfinder pipeline to extract all possible ortholog pairs between the two species. Among a total of 27,655 *A. thaliana* protein-coding gene models, 23,549 were paired with at least one *S. parvula* homolog. Similarly, 22,541 out of 24,769 *S. parvula* gene models were paired with at least one *A. thaliana* ortholog. The two reciprocal searches were merged, and redundant pairs were removed to generate 23,281 *S. parvula-A. thaliana* orthologous gene pairs (Table S2).

### Targeted quantification of hormones

For targeted analysis of hormones, samples were chromatographed on a 1.8 µm, 2.1 x 50 mm Agilent RRHD column. The mobile phase consisted of water containing 0.1% FA (solvent A) and ACN containing 0.1% FA (solvent B). The gradient was as follows at a flow rate of 0.3 mL/min: 0 min (5% B), 0-0.5 min (15% B), 0.5-2.5 min (40% B), 2.5-3.5 min (95% B), 3.5-4 min (95% B), 4-4.3 min (60% B), 4.3-5.5 (5% B) and 5.5-9.5 min (5% B) for re-equilibration. Quantification was performed using multiple reaction monitoring (MRM) in dynamic MRM mode. For each analyte, two MRM transitions were monitored: a primary transition (quantifier ion) used for quantitation and a secondary transition (qualifier ion) used for confirmation of analyte identity. Analyte identification was based on retention time matching with authentic standards and consistency of the quantifier-to-qualifier ion ratio. Collision energies voltages were optimized individually for each analyte using standard solutions. Deuterated compounds were used as internal standards to correct for extraction efficiency, ionization variability, and instrument drift. Data were acquired and processed using Agilent MassHunter-Qualitative and Quantitative Analysis software.

### Raman spectroscopy-based quantification for select metabolites

Raman spectra were acquired using a Renishaw inVia Reflex spectrometer equipped with a 785 nm laser, a 50X long-working-distance (LWD) air objective, and a 1200 lines/mm grating. Samples were mounted on stainless-steel slides to minimize background Raman signal. Spectra of the standards were collected in extended-scan mode over the 100–3200 cm⁻¹ range at 50% laser power (90 mW) with a 10 s exposure time. Raw spectra were preprocessed following previously reported protocols ^93–95^, including removal of cosmic-ray artifacts and subtraction of background signal, using WiRE 5.6 software. Metabolite standards for Cellulose (CAS 9004-34-6), Xylan (CAS 9014-63-5), L-Glutathione reduced (CAS 70-18-8), Methyl jasmonate (CAS 39924-52-2), Indole-3-acetic acid (IAA/Auxin; CAS 87-51-4), Abscisic acid (CAS 21293-29-8), L-Glutamine (CAS 56-85-9), L-Proline (CAS 147-85-3), L-Ascorbic acid (CAS 50-81-7), and D-(+)-Raffinose pentahydrate (CAS 17629-30-0) were purchased from Sigma-Aldrich and Cayman Chemical.

Metabolite scores were quantified by Direct Classical Least Squares (DCLS) regression against the curated metabolite reference library described above; the resulting coefficients were treated as semi-quantitative indicators of relative lipid abundance. DCLS, a supervised multivariate method grounded in the Beer–Lambert law, models the measured spectrum (D) as a linear combination of known reference spectra (S) weighted by concentration-related coefficients (C), plus residual error (E): D = C·Sᵀ + E.

### Data visualization and statistical tools used

Statistical analyses and data visualization were performed in R version 4.2.3 using RStudio version 2025.09.1+401. Specific sample sizes, treatment durations, statistical tests, and multiple-testing correction methods are indicated in the corresponding figure legends. Final figure assembly and formatting were performed using Microsoft PowerPoint for Mac.

## DATA AVAILABILITY

All sequence data generated in this study are available through the NCBI BioProject PRJNA1523682 under accession number SRA: GSE346137. All metabolite data and processed transcriptomic datasets used to construct the transcriptomic networks are provided in the Supplementary Tables.

## AUTHOR CONTRIBUTIONS

TTN conducted the main experiments. JRG, RSG, SW and HH contributed to physiological, biochemical, and transcriptomic data generation. MI and TTN performed the Raman spectroscopy experiments supervised by MRG and EGK conducted the time-lapse imaging experiments supervised by JRD. TTN, JRG, RSG, SW, and MD analyzed data. MD supervised and designed the overall experiment and data analysis. TTN, JRG, RSG, MI, and EGK contributed to writing Methods. TTN and MD interpreted the results and wrote the manuscript. JVM, MRG, JRG, RSG, SW, EGK, and JRD reviewed and provided critical feedback. MD, MRG, and JRD acquired funding and provided resources for the study.

## Acknowledgements

The authors thank Dr. Pradeep Kachroo and Dr. Keshun Yu from the Department of Plant Pathology, University of Kentucky, for customizing analytical LC-MS pipelines to quantify hormones from tissue samples. We also thank Dr. Rogério Gomes dos Santos, Advanced Microscopy and Analytical Core at LSU for assistance with confocal microscopy. We are grateful to Dr. Prava Adhikari Pantha, Anamaria Bao-Loc-Trung, Luke Guidry, Sylvie Bonner, Nithya Santhosh, and Laura L. Doan for their assistance in handling large volumes of tissue harvesting and sample preparation. We thank Dr. John Larkin for his constructive feedback. The authors also acknowledge the LSU High Performance Computing services for providing computational resources needed for data analyses. This work was supported by the US National Science Foundation (EDGE 1923589 to M.D. and CAREER 2045640 to M.R.G), and the Department of Energy (BER DE-SC0022985 to J.R.D and M.D.) awards. This work was inspired in part by discussions with and published work led by Dr. Hans Bohnert (1944–2025). MD gratefully acknowledges his enduring influence and his profound scientific contributions to the field.

## SUPPLEMENTARY FIGURES AND VIDEOS

**Figure S1.**
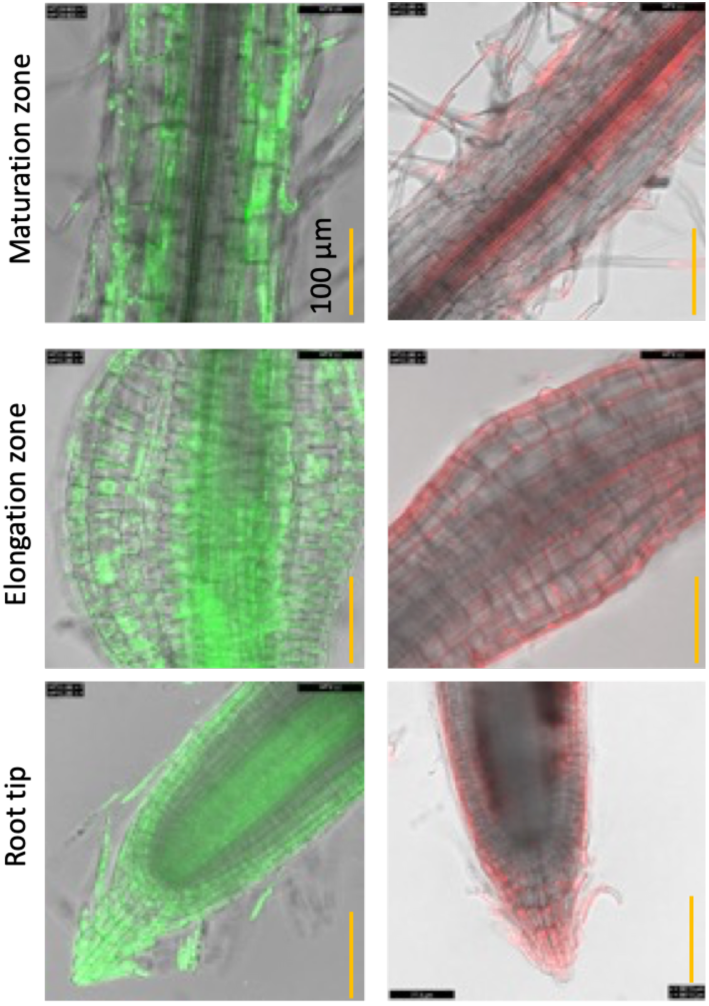
*S. parvula* roots stained with CoroNa Green (left column) and Propidium iodide (right column). 10-day-old seedlings were imaged after 3 days of exposure to 200 mM NaCl. Although sodium accumulated in cells across all developmental zones, most cells maintained intact plasma membranes that excluded PI.

**Figure S2.**
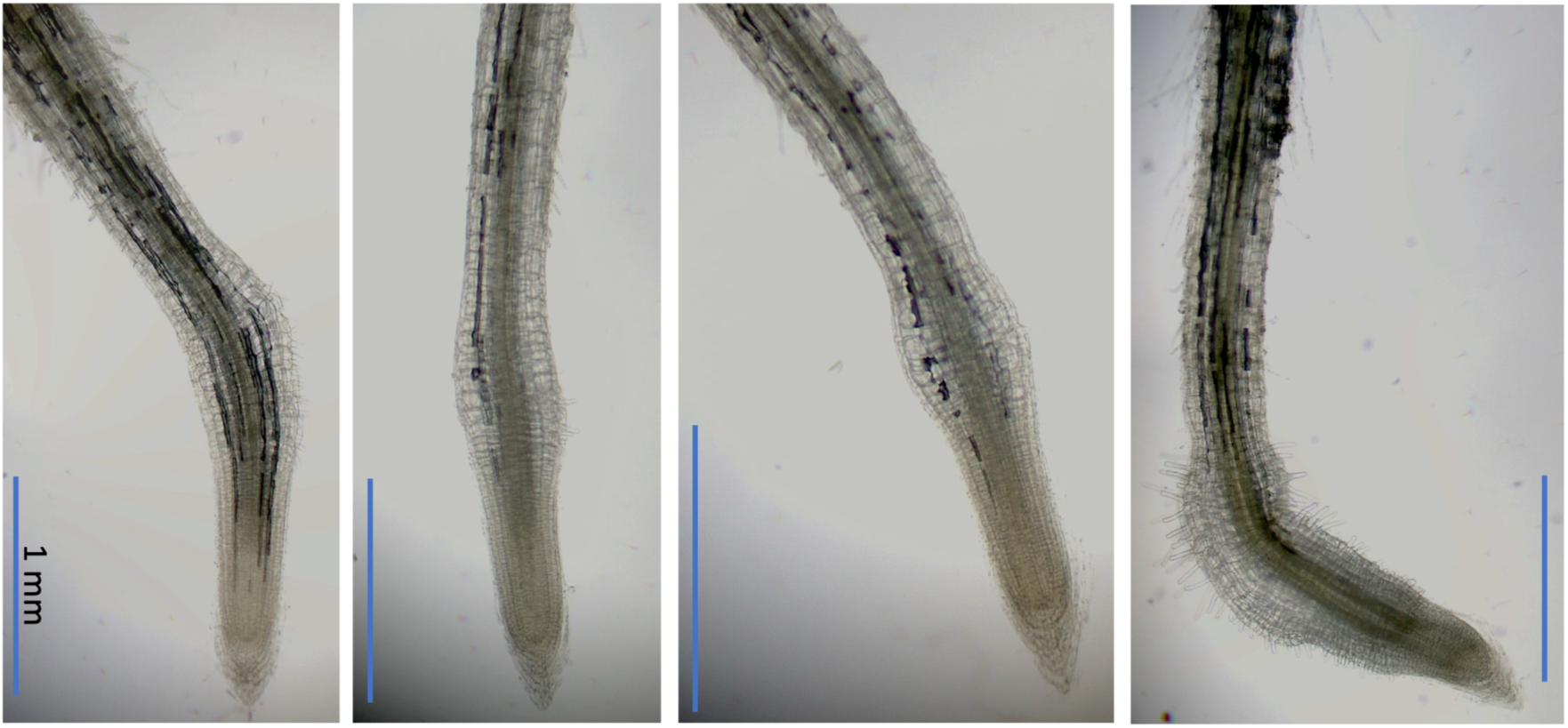
Morphological variation in the bulged zone of *S. parvula* seedlings after being treated with salt. 10-day-old seedlings were imaged after 3 days of exposure to 200 mM NaCl. Root hairs were either present or absent within the bulged region, and radial expansion could occur symmetrically or asymmetrically around the root axis.

**Figure S3.**
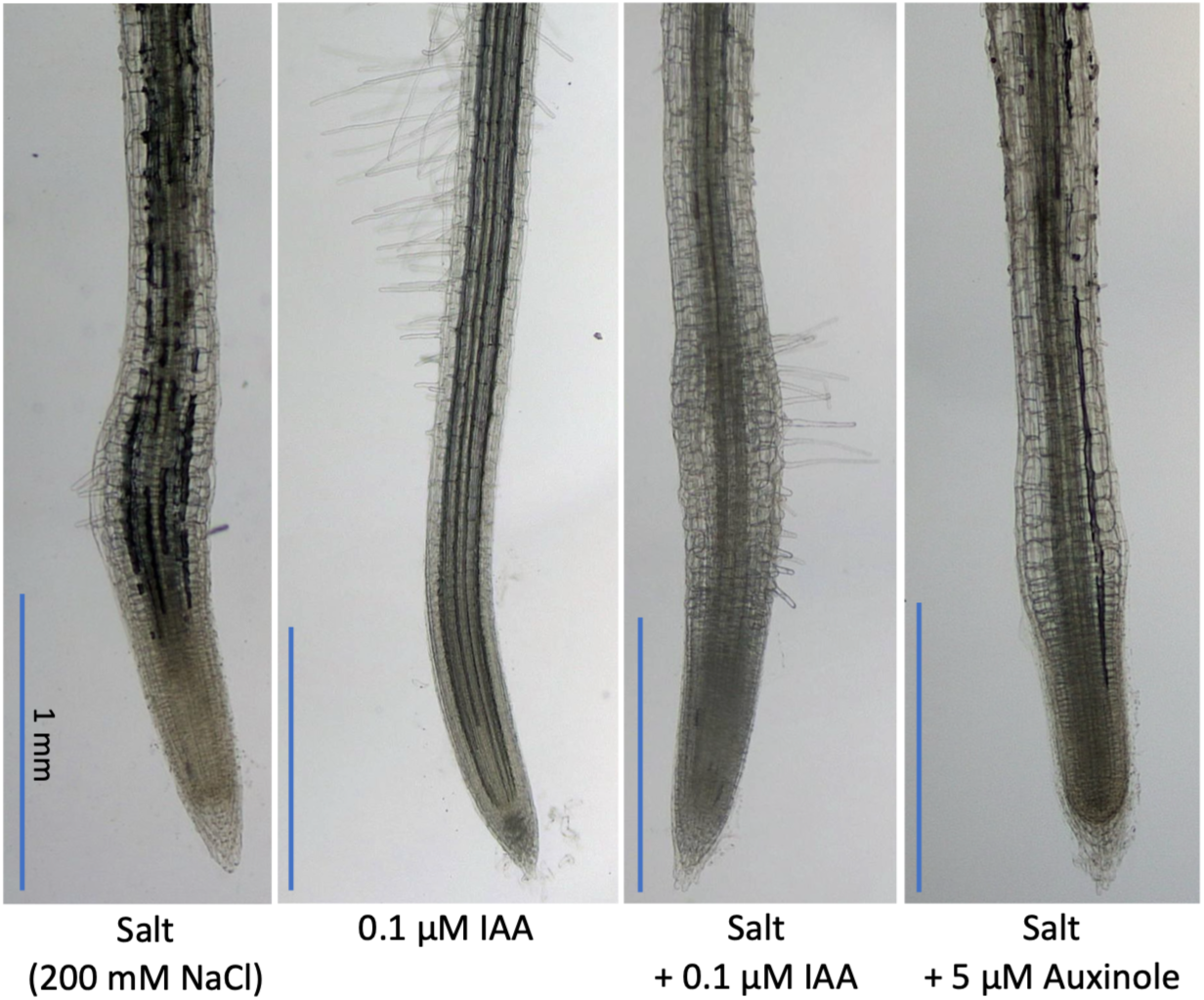
Auxin promotes root hair development within the bulged zone in salt-treated roots but is insufficient to induce bulged zone formation in the absence of salt in *S. parvula* primary roots. IAA is used as externally supplied auxin and auxinole is used as a synthetic antagonist that blocks endogenous auxin signaling. Images were taken after 3 days of treatment.

**Figure S4.**
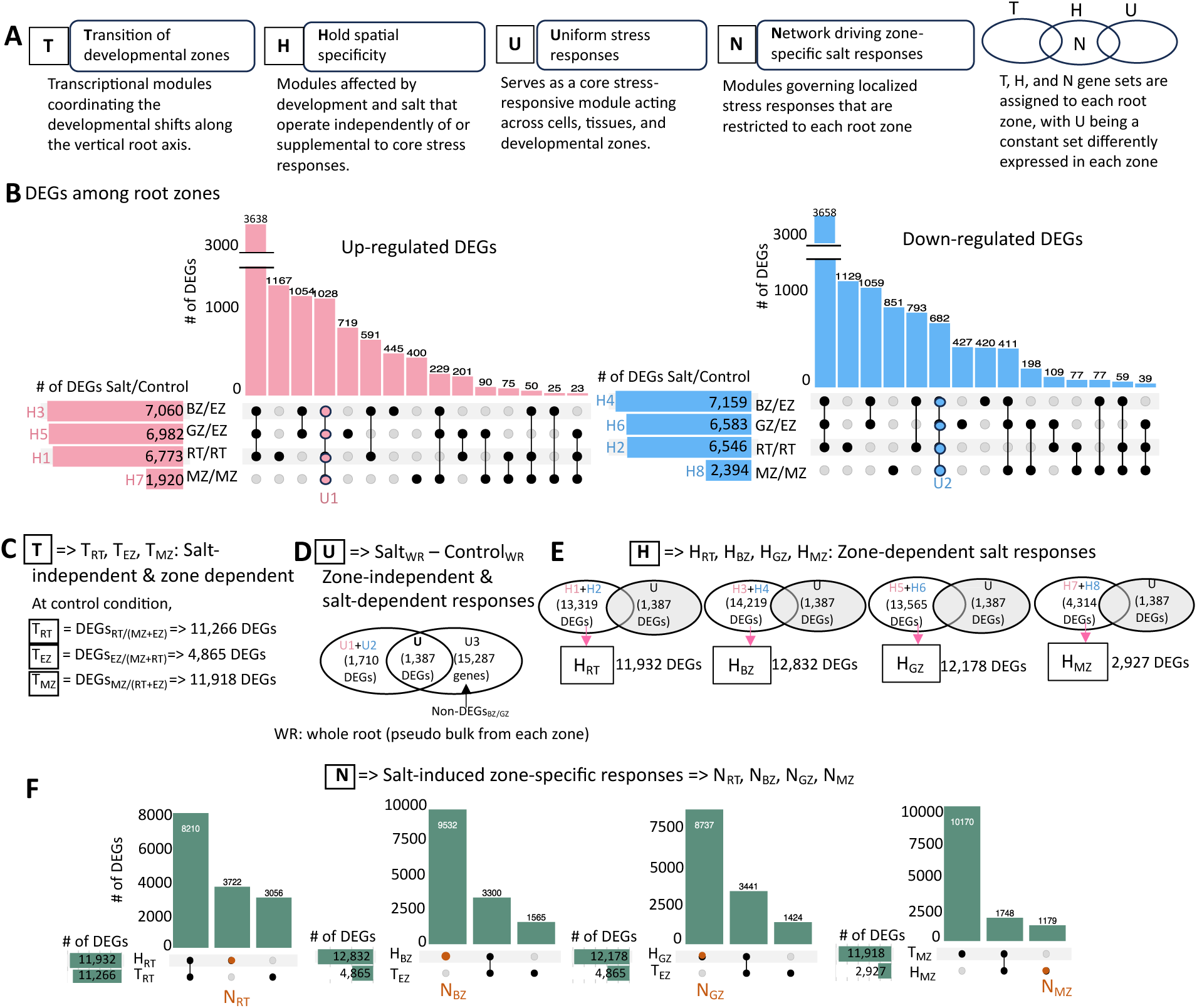
Conceptional dissection of total transcriptomes into T, H, U, and N modules. (A) The conceptual basis for the systematic dissection of transcriptional networks that add complexity to the baseline developmental networks under control conditions. (B) Differentially expressed genes across pairwise comparisons between salt-treated and control samples in different root zones. (C) Gene sets as defined by developmental DEGs from control conditions (T). **(D)** Differentially expressed genes in response to salt across all root zones representing generic whole-organ stress effects (U). **(E)** Genes differentially expressed within root zones but affected by salt (H). **(F)** Genes differentially expressed in response to salt within specific root zones, excluding those which are expressed entirely due to just developmental changes highlighting position-specific stress effects (N).

## SUPPLEMENTARY TABLES

**Table S1.** Raw read counts and normalized expression values, reported as TPM, for all sectioned *S. parvula* root samples under control and 200 mM NaCl treatments.

**Table S2.** List of orthologous gene pairs between *S. parvula* and *A. thaliana*.

**Table S3.** Lists of differentially expressed genes (DEGs) identified between root zones.

**Table S4.** Co-expressed genes grouped into five clusters and their functional enrichment results, corresponding to Fig. 4D.

**Table S5.** Lists of genes assigned to the T, H, U, and N sets, corresponding to Fig. 4E and S4.

**Table S6.** Lists of DEGs used to construct the T, H, U, and N gene-correlation networks, corresponding to Fig. 4F.

**Table S7.** Lists of DEGs assigned to functional processes represented in the N networks, corresponding to Fig. 4G.

**Table S8.** Functional enrichment results for DEGs shared between or unique to the N_BZ_ and N_GZ_ sets, corresponding to Fig. 5A. This table also includes the 129 DEGs with the highest expression levels in EZ, BZ, and GZ and their associated functional enrichment results, corresponding to Fig. 5A and 5B.

**Table S9.** Lists of hormone pathway-associated genes and their normalized expression values, reported as TPM, corresponding to Fig. 6.

**Table S10.** Lists of DEGs associated with salt or osmotic stress, ROS response, and cell death in the bulged and gap zones, corresponding to Fig. 7D, 8E, and 8F.

## SUPPLEMENTARY FILES

**Supplementary Video S1. Time-lapse imaging of *S. parvula* primary root growth under control conditions.** Primary root continues to grow without a bulge forming. Imaging began immediately after treatment and continued for 16 hours.

**Supplementary Video S2. Time-lapse imaging of salt-induced root growth reprogramming at 200 mM NaCl.** Salt stress transiently pauses primary root growth, induces a bulge formation, and is followed by resumed root elongation. Imaging began 2 hours after treatment and continued for 16 hours.

**Supplementary Video S3. Time-lapse imaging shows primary root growth inhibition in *S. parvula* under 250 mM NaCl. Primary root growth was inhibited and no bulge was formed.** Imaging began 30 minutes after treatment and continued for 16 hours.

**Supplementary Video S4. Time-lapse imaging shows primary root growth in *S. parvula* treated with ABA.** No bulge or gap were formed. Imaging began immediately after treatment and continued for 16 hours.

